# Age-related decline in cortical inhibitory tone strengthens motor memory

**DOI:** 10.1101/2020.12.03.410308

**Authors:** Pierre Petitet, Gershon Spitz, Uzay E. Emir, Heidi Johansen-Berg, Jacinta O’Shea

**Affiliations:** Wellcome Centre for Integrative Neuroimaging, FMRIB Centre, Nuffield Department of Clinical Neurosciences, Oxford, United Kingdom; Centre de Recherche en Neurosciences de Lyon, Equipe Trajectoires, Inserm UMR-S 1028, CNRS UMR 5292, Université Lyon 1, Bron, France; Turner Institute for Brain and Mental Health, Monash University, Melbourne, Australia; School of Health Sciences, Purdue University, West Lafayette, Indiana, USa; Weldon School of Biomedical Engineering, Purdue University, West Lafayette, Indiana, USA; Wellcome Centre for Integrative Neuroimaging, Oxford Centre for Human Brain Activity (OHBA),University of Oxford Department of Psychiatry, Warneford Hospital, Warneford Lane, Oxford, United Kingdom

**Author notes:** Contributed equally to this work (Pierre Petitet, Gershon Spitz).

## Abstract

Ageing disrupts the finely tuned excitation/inhibition balance (E:I) across cortex, driven by a natural decline in inhibitory tone (*γ*-amino butyric acid, GABA). This causes functional decrements. However, in young adults, experimentally lowering GABA in sensorimotor cortex enhances adaptation memory. Therefore, using a cross-sectional design, here we tested the hypothesis that as sensorimotor cortical GABA declines naturally with age, adaptation memory would increase, and the former would explain the latter. Results confirmed this prediction. To probe causality, we used brain stimulation to further lower sensorimotor cortical GABA during adaptation. Across individuals, how stimulation changed memory depended on sensorimotor cortical E:I. In those with low E:I, stimulation increased memory; in those with high E:I stimulation reduced memory. Thus, we identify a form of motor memory that improves naturally with age, depends causally on sensorimotor cortex neurochemistry, and may be a potent target for motor skill preservation strategies in healthy ageing and neurorehabilitation.

## Introduction

Motor capacities decline with age^1, 2^. As the brain and body become older, movements lose speed^3, 4^, strength^5^ and coordination^6^. This natural loss of function is exacerbated by motor disorders which rise sharply with age (e.g., stroke, sarcopenia, Parkinsonism). As the elderly population increases^7^, there is a need for strategies to counteract and compensate for age-related motor decline.

During ageing, the motor system must adapt continuously to ongoing neuro-musculo-skeletal change. Brain plasticity enables this. Plasticity is essential to learn new motor skills, adapt and retain existing ones, and to rehabilitate functions impaired by disease^8, 9^. Thus plasticity plays an important role in mitigating age-related motor decline^10, 11^.

Unfortunately, plasticity also declines with age^12^, especially in the motor domain^13–15^. A major cause is the dysregulation of the finely tuned balance between cortical excitation and inhibition (E:I)^10^. Across cortex, E:I is disrupted because *γ*-aminobutyric acid (GABA), the major inhibitory neurotransmitter, declines with age, both in animals^16, 17^ and humans^15, 18–26^. Regional decline of cortical GABA causes a loss of inhibitory tone, and this is associated with decrements in functions localized to the affected regions^27–29^. For example, in somatosensory cortex lower GABA (i.e., higher E:I) is associated with poorer tactile discrimination, both in young and old adults ^20, 30^. In primary motor cortex (M1), age-related decline of inhibitory tone is associated with poorer upper-limb dexterity^23^, postural imbalance^31, 32^, and impaired ability to suppress automatic responses^26^.

By contrast, here we tested the hypothesis that, as M1 GABA declines with age, a specific form of upper limb motor function – adaptation memory – would *increase*. Across the lifespan, adaptation is that property of the motor system that enables individuals to counteract perturbations by adjusting their movements and thus maintain successful motor performance ^33, 34^. There is a wealth of evidence that adaptation is preserved (or somewhat impaired) in healthy ageing^35–46^. After a perturbation is removed, adaptation memory is expressed as an *after-effect* – a movement bias in the direction opposite the perturbation. The strength of adaptation memory is indexed by the persistence over time of this after-effect. In our previous work, in young adults, we showed that experimentally lowering M1 inhibitory tone during adaptation increases persistence of the after-effect^47, 48^. Here, we reasoned that if after-effect retention depends *causally* on M1 inhibitory tone, then owing to age-related M1 GABA decline, this form of memory would *increase naturally with age*.

This hypothesis was confirmed in a cross-sectional study of thirty-two healthy older adults (mean age: 67.46 years, *s.d.*: 8.07). Using magnetic resonance spectroscopy to quantify neurochemistry, we showed that M1 GABA declines with age. Using prism adaptation^49^, we showed that retention increases with age. A mediation analysis subsequently confirmed that as GABA declines with age, adaptation memory increases, and the former explains the latter. To demonstrate causality, we intervened experimentally with excitatory anodal transcranial direct current stimulation - to try and further lower M1 GABA^50–52^ and thus further increase adaptation memory. On average, stimulation did not increase memory in this age group. Rather, a moderation analysis showed that how stimulation changed memory depended on individuals’ motor cortical E:I. Stimulation increased retention in individuals with low E:I, but decreased retention in individuals with high E:I.

In summary, we identified a specific domain of motor functional plasticity that improves with age, as a natural consequence of motor cortical inhibitory decline. This memory function can be further enhanced by neurostimulation, but only in individuals least affected by age-related dysregulation of motor cortical E:I. These findings challenge the prevailing view of ageing as inevitable functional decline. Whereas learning of new motor skills may decline, the capacity to maintain adaptation of existing skills improves naturally with age. That adaptation memory is enhanced naturally with age indicates it may have untapped potential as a target for training strategies that aim to preserve, improve or restore motor function in healthy or pathological ageing^48^.

## Results

### Retention increases with age

First we tested the prediction that adaptation memory increases with age. We used a cross-sectional correlational design to measure the continuous effect of ageing across a mid- to late-life sample. This avoids the confounds inherent in a between-groups “young vs. old” design caused by gross differences in body, brain and behaviour. In Experiment 1 thirty two healthy male volunteers aged between 49 and 81 (mean age: 67.46 years, *s.d.*: 8.07; Table S1) performed a session of prism adaptation (PA) with their dominant right hand. Only men were recruited to avoid the additional variability caused by the impact of ovarian hormone fluctuations on neurotransmitter concentration in women^53–55^ (see *Methods*).

The behavioural protocol was similar to previous work from our laboratory^48, 56^ (full details in *Methods*). Following adaptation, retention of the after-effect was assessed after a short (10 minutes) and long (24 hours) interval (Fig. S1). Effects were analysed statistically using linear mixed-effect models (LMMs) with maximal random structure. This allowed us to assess both the average lateral error across task blocks and the stability of the error within blocks, while controlling for random effects of inter-individual variation.

Fig.1a shows the pointing error data, plotted as changes from baseline (pre-adaptation) accuracy. Throughout adaptation, participants made rapid pointing movements at a 10° left and right target, while wearing prism glasses that displaced their visual field 10° to the right. During prism exposure (Blocks E1-6) participants gradually corrected their errors. The learning and forgetting dynamics are visible within and across blocks. At prism onset participants exhibited a large rightward error (Fig.1a; Block E1, trial 1: mean 7.77°, s.e.m.: 1.05°, one-sample t-test compared to zero: *t*_(31)_ = 7.43*, p <* 0.001, Cohen’s *d* = 1.31) which was corrected gradually across trials and blocks (E1-6) until performance stabilized (E6) close to restored baseline accuracy (main effect of Trial within Block: *t*_(3185)_ = −11.28*, p <* 0.001, 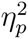 = 0.47; main effect of Block: *t*_(3185)_ = −9.05*, p <* 0.001, 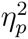 = 0.73; Table S3 - model 1).

As participants adapted gradually to the rightward visual shift, a consequent leftward after effect developed, measured in interleaved blocks, critically without prisms and without visual feedback (Fig. 1a; Blocks AE1-6; mean normalised error: −6.66°, *t*_(2865)_ = −16.94, *p <* 0.001, 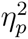 = 0.90; Table S3 - model 2). This prism after-effect (AE) is the key experimental measure. On AE trials, the absence of visual feedback prevents error-based learning and requires participants to rely on internal representations of sensed limb position to guide their movements. Thus, the leftward AE expresses the visuomotor transformation acquired during prism exposure. Its persistence after prism removal is the measure of adaptation memory. The AE was measured after each block of prism exposure (AE1-6, Fig. S1). Initially memory was labile: on the first trial of the first block the AE was large (−6.99°), but across the 15 trials of the first block it decayed by 2.70° on average. Subsequent blocks of prism exposure led the AE to gradually stabilize, evidenced by the progressive flattening of slopes across blocks AE1-6 (interaction Trial × Block: *t*_(2865)_ = −3.33, *p* = 0.001, 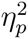 = 0.26; Fig. 1a; Table S3 - model 2). Thus, our protocol induced an adaptation memory trace that consolidated gradually across the Adaptation phase.

**Figure 1:**
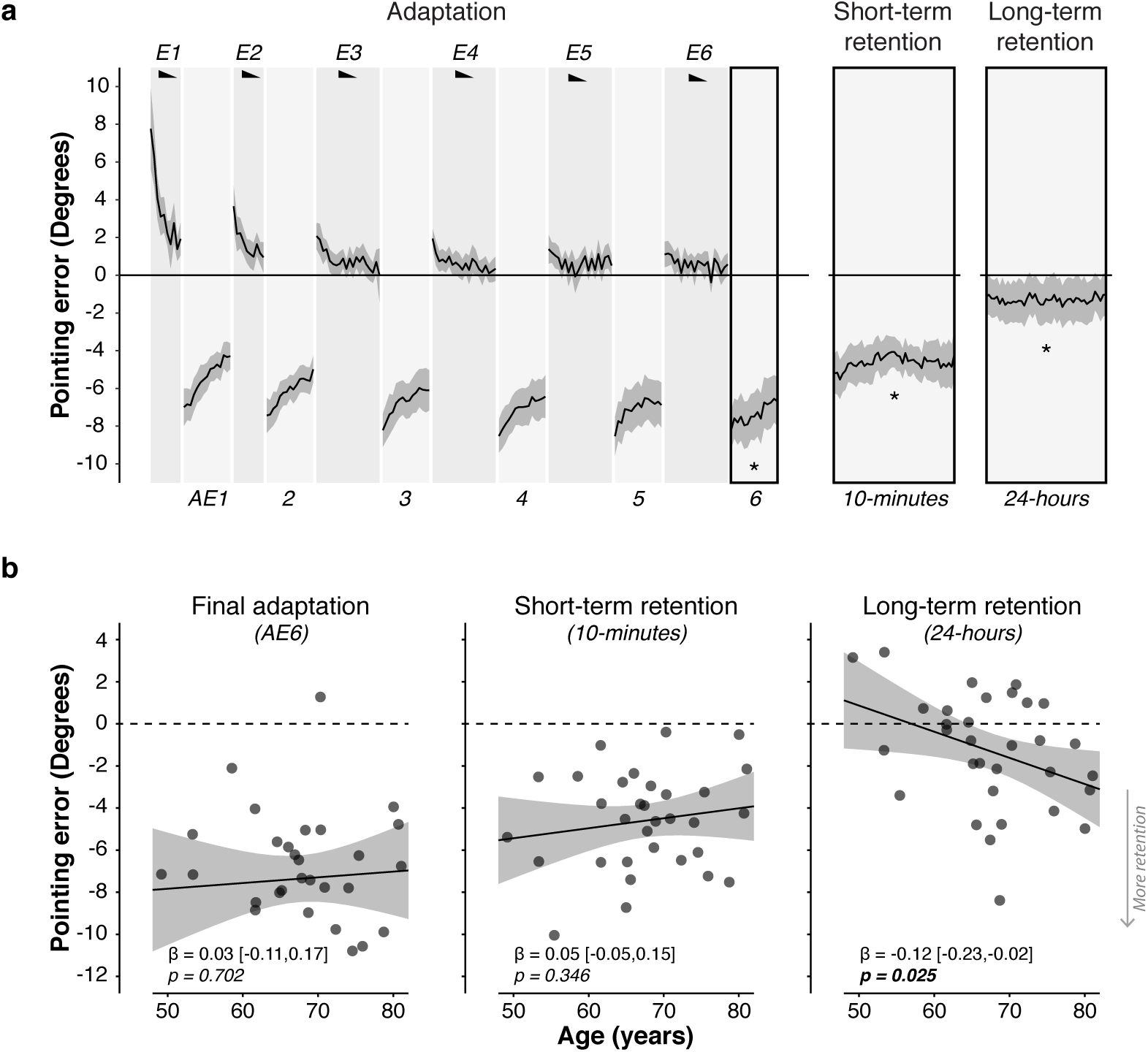
Long-term retention of prism adaptation is higher in older adults. **a.** Group mean pointing errors (±1 s.e.m.) expressed as change from baseline accuracy (y=0). Positive y-axis values are rightward errors, negative leftward. Black wedges indicate blocks in which prisms were worn. During right-shifting prism exposure (E1-6), visual feedback enabled participants to correct their rightward pointing errors across trials. Consequent leftward after-effects were measured in intervening blocks without visual feedback throughout adaptation (AE1-6). After-effect retention was measured post-adaptation after a short (10 minutes) and long (24 hours) interval. There was significant retention at both time points. **b.** Age had no effect on the after-effect magnitude acquired by the end of adaptation (block AE6), nor on short-term retention (10 minutes). The key finding was that older adults showed significantly greater long-term retention (24-hours). Full statistics are in Tables S3 & S4.

The critical measure of memory was AE retention post-adaptation (Fig. 1a-b). After 10 minutes of blindfolded rest there was significant short-term retention (mean error: −4.61°, s.e.m.: 0.41°, *t*_(1434)_ = −11.36, *p <* 0.001; one sample t-test of mean retention: *t*_(31)_ = −11.18, *p <* 0.001, Cohen’s *d* = -1.98; Table S3 - model 3). Long-term retention, measured 24 hours later, was also significant (mean error: −1.30°, s.e.m.: 0.48°, *t*_(1434)_ = −2.75, *p* = 0.006, 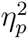 = 0.19; one sample t-test of mean retention: *t*_(31)_ = −2.70, *p* = 0.01, Cohen’s *d* = -0.48; Table S3 - model 4). In our previous work with young adults long-term retention was not significant^48^. The AE was stable at both time points, indicated by no change in error across trials (main effect of Trial: both *p >* 0.38).

Our hypothesis was that AE retention would increase with age. Fig. 1b plots the results. Age had no effect on the AE magnitude acquired by the end of prism exposure (Block AE6), nor on short-term retention (both *p >* 0.35; Fig. 1b; Table S4 - models 1 & 2). However, older age was associated with greater long-term retention (Age × AE_24hrs_: *t*_(1432)_ = −2.24*, p* = 0.025, 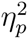 = 0.14, Fig. 1b, Table S4 - model 3). This is the key finding. This association remained significant when controlling for the AE at the two preceding time points (AE6 and 10-min retention), and when controlling for average reaching speed during prism exposure (slower movements, expected in ageing, could arguably favour retention; Table S4 - models 4-6).

### Motor cortical inhibitory tone declines with age

Next we tested for an expected decrease in motor cortical inhibitory tone with age. Three Tesla magnetic resonance spectroscopy was used to quantify neurochemical concentration in left sensorimotor cortex (labelled “M1”), and in a control region of occipital cortex (labelled “V1”; see *Methods*; Fig. S2). The metabolites of interest were GABA and Glutamix (”Glx”= Glutamate + Glutamine, since these two metabolites cannot be distinguished reliably at 3 Tesla). As expected, in both regions, age was associated with significant grey matter atrophy (both *p <* 0.002), which could indirectly lower neurochemical concentration estimates. Hence, all analyses of neurochemistry ruled out this potential confound by controlling for grey and white matter fractions within each region (see *Methods*). To minimize multiple comparisons, analyses focused on the ratio of excitation:inhibition (E:I = Glx:GABA). If an effect was significant, follow-up analyses assessed the individual contributions of Glx and GABA.

Figure 2 shows the results. Multiple linear regressions showed that sensorimotor cortex E:I increased with age (standardised *β_age_* = 0.66, *t*_(18)_ = 2.09, *p* = 0.051, 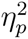 = 0.10; Table S5 - model 1). As predicted, across individuals, as age increased, M1 GABA concentration decreased (standardised *β_age_* = −0.74, *t*_(17)_ = −2.48, *p* = 0.024, 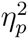 = 0.14; Table S5 - model 2). There was no such relationship with Glx (standardised *β_age_* = −0.23, *t*_(17)_ = −0.68, *p* = 0.51, 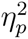 = 0.02;

**Figure 2:**
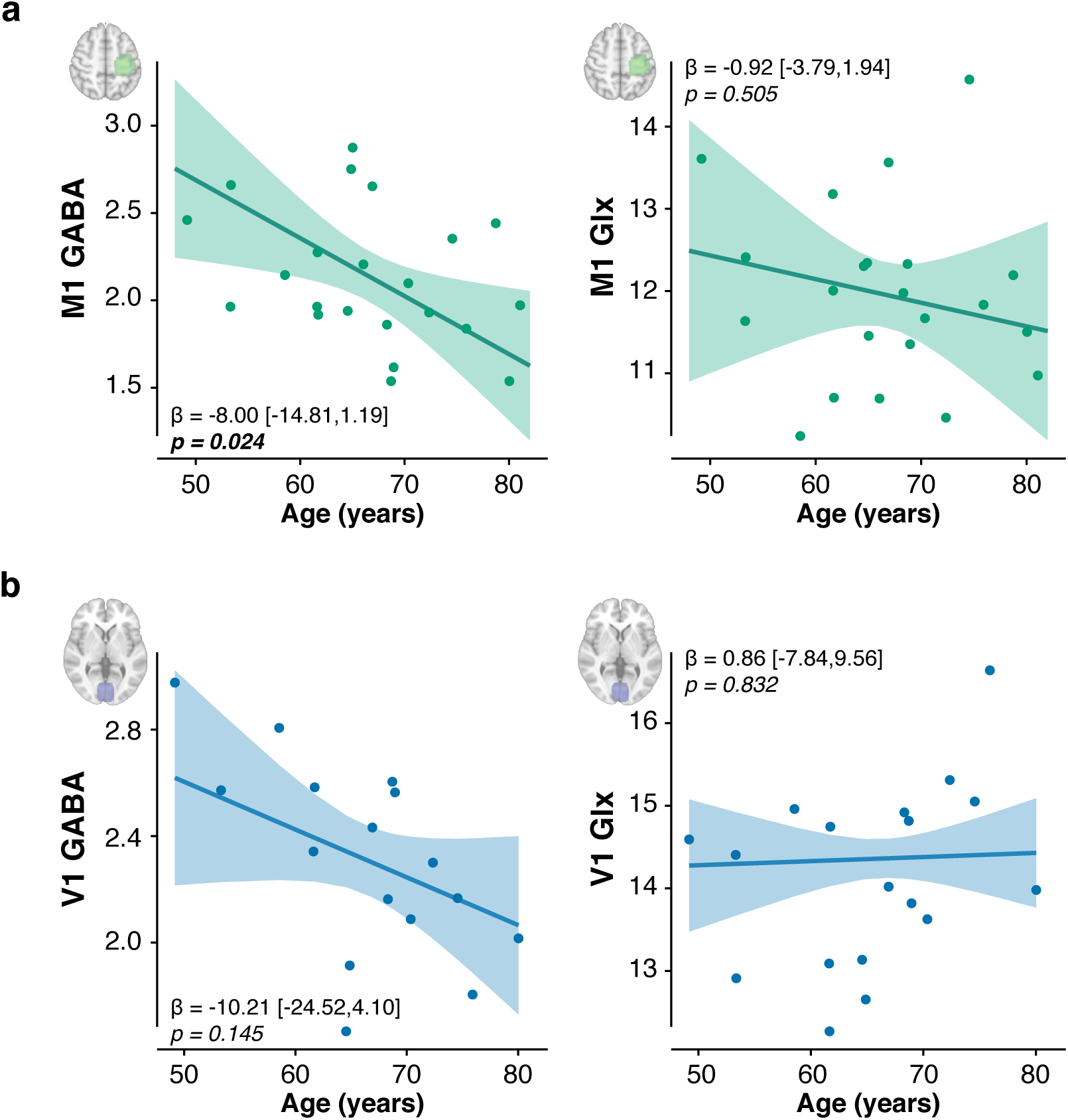
Motor cortical inhibitory tone is lower in older adults. **a.** The concentration of GABA but not Glutamix (Glutamate + Glutamine, Glx) was associated negatively with age in the left sensorimotor cortex (labelled “M1”). **b.** There was no significant association between age and neurochemical concentration in occipital cortex (labelled “V1”). For each voxel and neurotransmitter, plotted relationships control for the fraction of grey matter and white matter, and the other neurotransmitter. Absolute concentrations are expressed in arbitrary units. Full statistical details are in Table S5.

Fig. 2a, Table S5 - model 3).

In the anatomical control region (occipital cortex), there was a qualitatively similar pattern of age-related inhibitory decline, consistent with previous reports ^28, 29^. However this was not statistically significant, likely reflecting the impact of quality controls that reduced the size of the occipital dataset and consequently reduced power (Table S1 & S2). There was no significant relationship between neurochemistry and age in V1, not for E:I (standardised *β_age_* = 0.39, *t*_(12)_ = 1.46, *p* = 0.171, 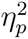 = 0.46), GABA (standardised *β_age_* = −0.40, *t*_(11)_ = −1.57, *p* = 0.145, 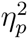 = 0.51) or Glx (standardised *β_age_* = 0.04, *t*_(11)_ = 0.22, *p* = 0.832, 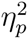 = 0.20; ; Table S5 - models 4-6).

### Lower motor cortical inhibitory tone is associated with greater long-term retention

Based on our previous work^48^, we hypothesized that lower motor cortical inhibitory tone would be associated with greater retention. Results confirmed this prediction (Fig. 3). Across individuals, higher sensorimotor cortex E:I was associated with a larger prism AE at retention 24-hours after adaptation (*t*_(980)_ = −5.40, *p <* 0.001, 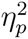 = 0.58; Table S6 - model 1). This relationship was driven by GABA: individuals with lower M1 GABA concentration showed greater retention (*t*_(978)_ = 5.04, *p <* 0.001, 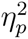 = 0.55; Fig. 3a, Table S6 - model 2). There was no such relationship with M1 Glx (*t*_(978)_ = 0.01*, p* = 0.99, 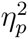 *<* 0.001; Fig. 3a, Table S6 - model 2). Thus, this memory effect was neurochemically specific (M1 GABA vs. M1 Glx: *z* = 3.56, *p <* 0.001). It was also anatomically specific (M1 GABA vs. V1 GABA: *z* = 2.80, *p* = 0.005): there was no relationship between retention and V1 metabolites - not for GABA, Glx or E:I (all *p >* 0.25; Fig. 3b, Table S6 - models 5 & 6). As before, the results were unchanged when controlling for average movement time during prism exposure (S6 - models 3, 4, 7, 8).

**Figure 3:**
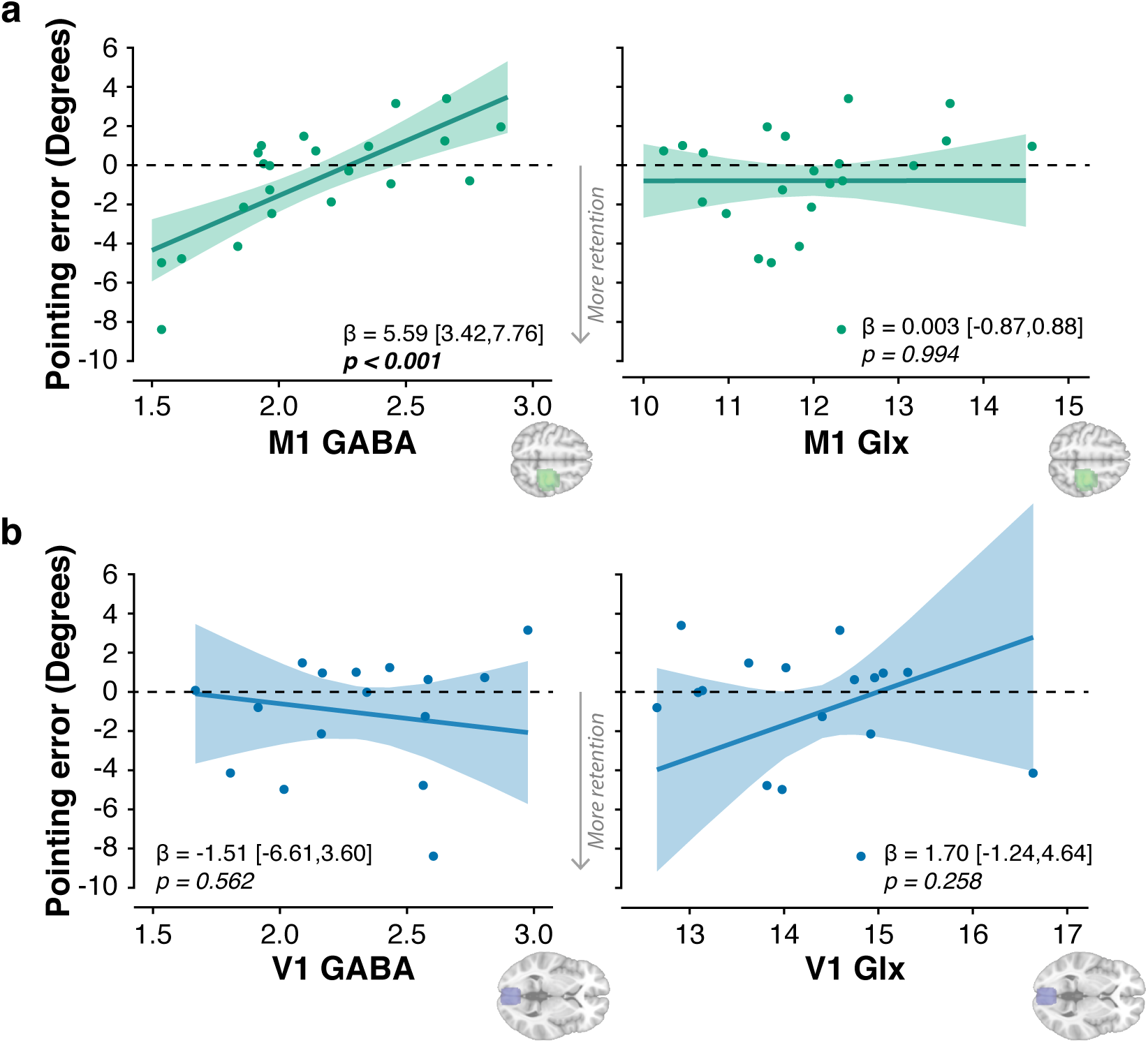
Lower motor cortical inhibitory tone is associated with greater long-term retention. Plot shows relationships between brain chemistry and the magnitude of prism after-effect retained 24 hours after adaptation. Negative values on the y-axis indicate retention. **a. Sensorimotor cortex (“M1”)** Across individuals, lower GABA was associated with greater retention. There was no relationship with Glx (Glutamate + Glutamine). **b. Occipital cortex (“V1”)** There was no relationship between GABA or Glx and 24-hour retention. For each voxel and neurotransmitter, relationships control for the fraction of grey matter and white matter, and the other neurotransmitter. Absolute concentrations are expressed in arbitrary units. Full statistics details are in Table S6.

### Retention increases with age as a function of motor cortical inhibitory decline

Our key prediction was that as M1 GABA concentration declines with age, adaptation memory would increase, and the former would explain the latter. We used mediation analysis to formally test this hypothesis. Mediation analysis is well suited to a situation in which the independent variable (Age) may not directly influence the dependent variable (Long-term retention), but is instead hypothesized to do so indirectly via its influence on candidate mediators (M1 E:I, GABA, Glx). The extent to which the relationship between the independent and dependent variable is influenced by a mediator is termed the indirect effect. We tested the significance of indirect effects using a bootstrap estimation approach with 10,000 samples (see *Methods*).

Figure 4 shows that, as hypothesized, the effect of age on long-term retention was mediated by motor cortical E:I (*ab*_1_ = *−*0.41, 95%CI: [*−*1.45*, −*0.08], *p* = 0.017). More specifically, the indirect effect was driven by M1 GABA and not Glx. M1 GABA was a significant mediator (*ab*_1_ = *−*0.50, 95%CI: [*−*1.46*, −*0.16], *p* = 0.0086), accounting for 64% of the variance between age and long-term retention (Fig. 4, Table S7), while M1 Glx showed no such effect (*ab*_2_ = 0.018, 95%CI: [*−*0.095, 0.31], *p* = 0.74). When M1 neurochemistry was controlled for, age was no longer a significant predictor of 24-hour retention (*c′* = −0.28, *p* = 0.38), consistent with full mediation. Thus, age-related decline in sensorimotor GABA explains stronger adaptation memory in older age. Once again, results were unchanged when controlling for average movement time during prism exposure (Table S8).

**Figure 4:**
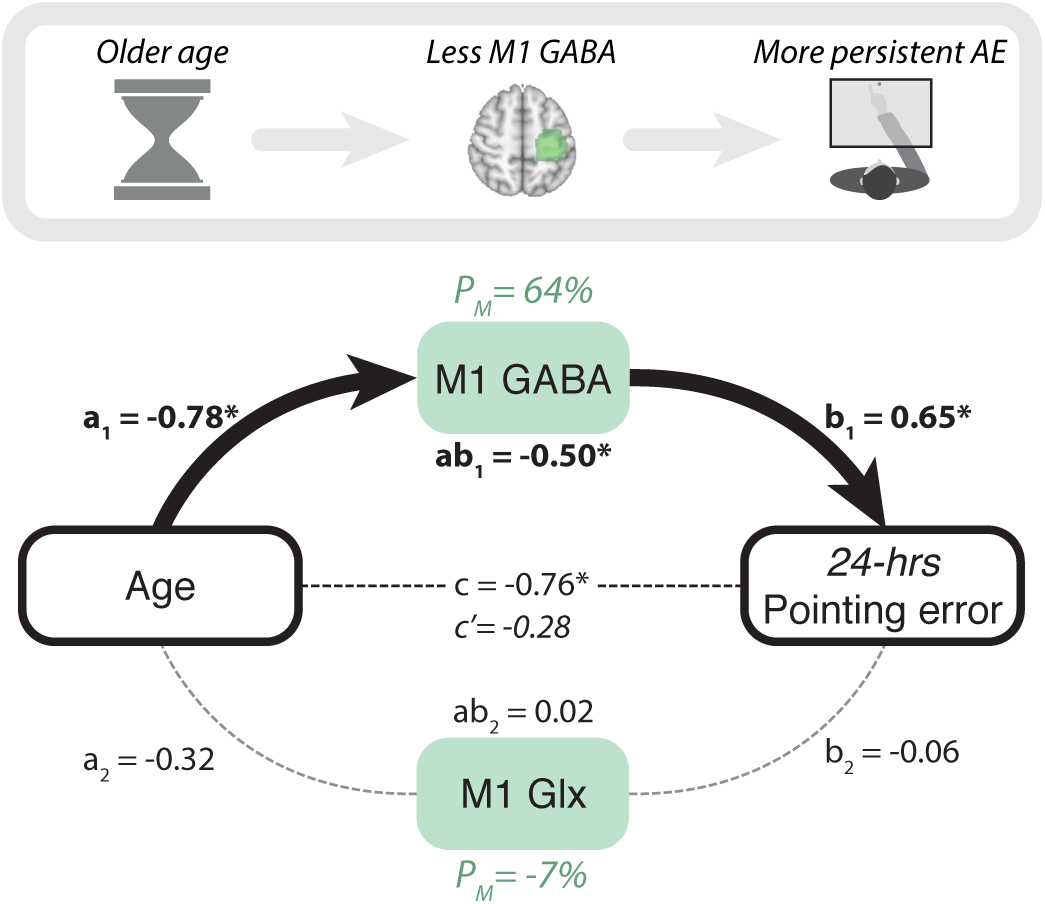
Adaptation memory is stronger in older age owing to the decline in motor cortical inhibitory tone. A mediation model tested whether M1 neurochemistry explained the relationship between age and retention. Consistent with our mechanistic hypothesis, GABA, but not Glx, mediated the positive relationship between age and 24-hour retention, explaining 64% of the variance. Standardised regression coefficients are reported next to the corresponding paths. Asterisks indicate significance (*p <* 0.05). Full statistics: Table S7. Independent variable: Age. Dependent variable: AE 24-hours post-adaptation. Mediators: M1 GABA and Glx (controlling for grey and white matter tissue fractions).

### How stimulation changes memory depends on motor cortical E:I

The mediation model indicated that the M1 GABA decline was responsible for the memory increase in older adults. However, the cross-sectional study design precludes direct causal inference^57^. Hence, to more directly test causation, we intervened experimentally with anodal transcranial direct current stimulation (a-tDCS). M1 a-tDCS has been shown to increase motor cortical E:I in young^50, 51, 58^ and older^52^ adults. In addition, we previously showed (in young adults) that M1 a-tDCS during adaptation increased short- and long-term retention, in proportion to the stimulation-induced E:I increase ^48^. However, given our finding in Experiment 1 (Fig. 2) – that M1 E:I is already naturally high in older age – we expected there may be a ceiling effect on further increasing E:I in some individuals. Thus, we expected that stimulation would, on average, be consequently less effective at increasing retention in older adults. Hence we tested the hypothesis that, across individuals, the magnitude of memory change induced by stimulation would depend causally on individuals’ intrinsic M1 E:I.

To test this hypothesis, a sub-set of twenty-five participants from Experiment 1 (mean age: 69.6 years, *s.d.*: 8.4; Table S1) consented to undergo a follow-up study (Experiment 2), in which tDCS (anodal/sham, counterbalanced repeated measures design) was applied in two weekly test sessions to left M1 during adaptation, and retention was assessed after 10 minutes and 24 hours (see *Methods*, Fig. S1).

Figure 5 shows the results for the group average. Stimulation had no effect on short-term retention (*t*_(2235)_ = 0.22, *p* = 0.83, 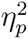 = 0.002; paired-samples t-test of mean retention, anodal vs sham: *t*_(24)_ = 0.21, *p* = 0.83, Cohen’s *d* = 0.04; Table S9 - model 1). Although long-term retention increased numerically, this was not significant (*t*_(2235)_ = −1.35, *p* = 0.18, 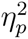 = 0.07; paired-samples t-test of mean retention, anodal vs sham: *t*_(24)_ = −1.32, *p* = 0.20, Cohen’s *d* = -0.26; Table S9 - model 4). The lack of a significant memory gain from stimulation across the group contrasts with our previous findings in young adults^48^.

**Figure 5:**
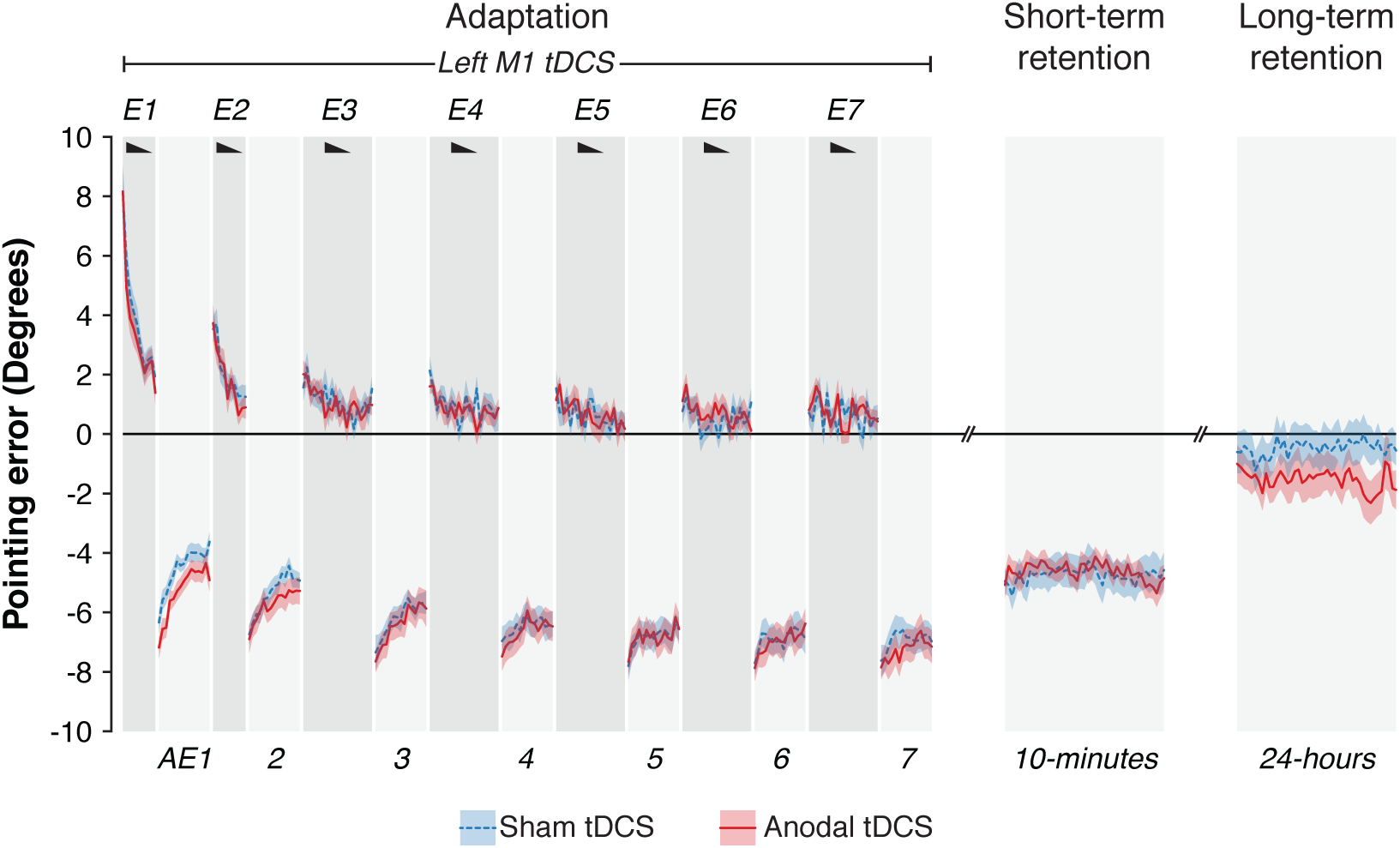
On average across older adults excitatory stimulation of M1 during adaptation did not increase retention. Timecourse of pointing errors for the same behavioural paradigm and graph conventions as in Fig. 1, except that stimulation (anodal or sham tDCS) was applied to left M1 throughout the adaptation phase. Errors are normalised to baseline (pre-adaptation) accuracy. Negative values on the y-axis indicate prism after-effects. After excitatory stimulation of M1 during adaptation, retention increased numerically but not significantly, contrary to our previous findings in young adults, but consistent with our expectations in this cohort of older adults.

To test the key hypothesis, that individual differences in intrinsic motor cortical E:I would influence the magnitude of stimulation-induced memory change, we conducted a moderation analysis. For all participants who had undergone an MRS scan in Experiment 1 (*n* = 16, Table S1) we added their M1 Glx:GABA levels from that scan to the linear mixed model analyses of the effect of stimulation on retention. As predicted, the effect of stimulation on long-term retention interacted significantly with motor cortical E:I (E:I × a-tDCS: *t*_(1419)_ = 2.40, *p* = 0.009, one-tail, 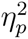= 0.26; Fig. 6; Table S9 - model 5). A similar pattern was observed for short-term retention (*t*_(1419)_ = 1.86, *p* = 0.003, one-tail, 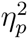 = 0.18; Table S9 - model 2).

**Figure 6:**
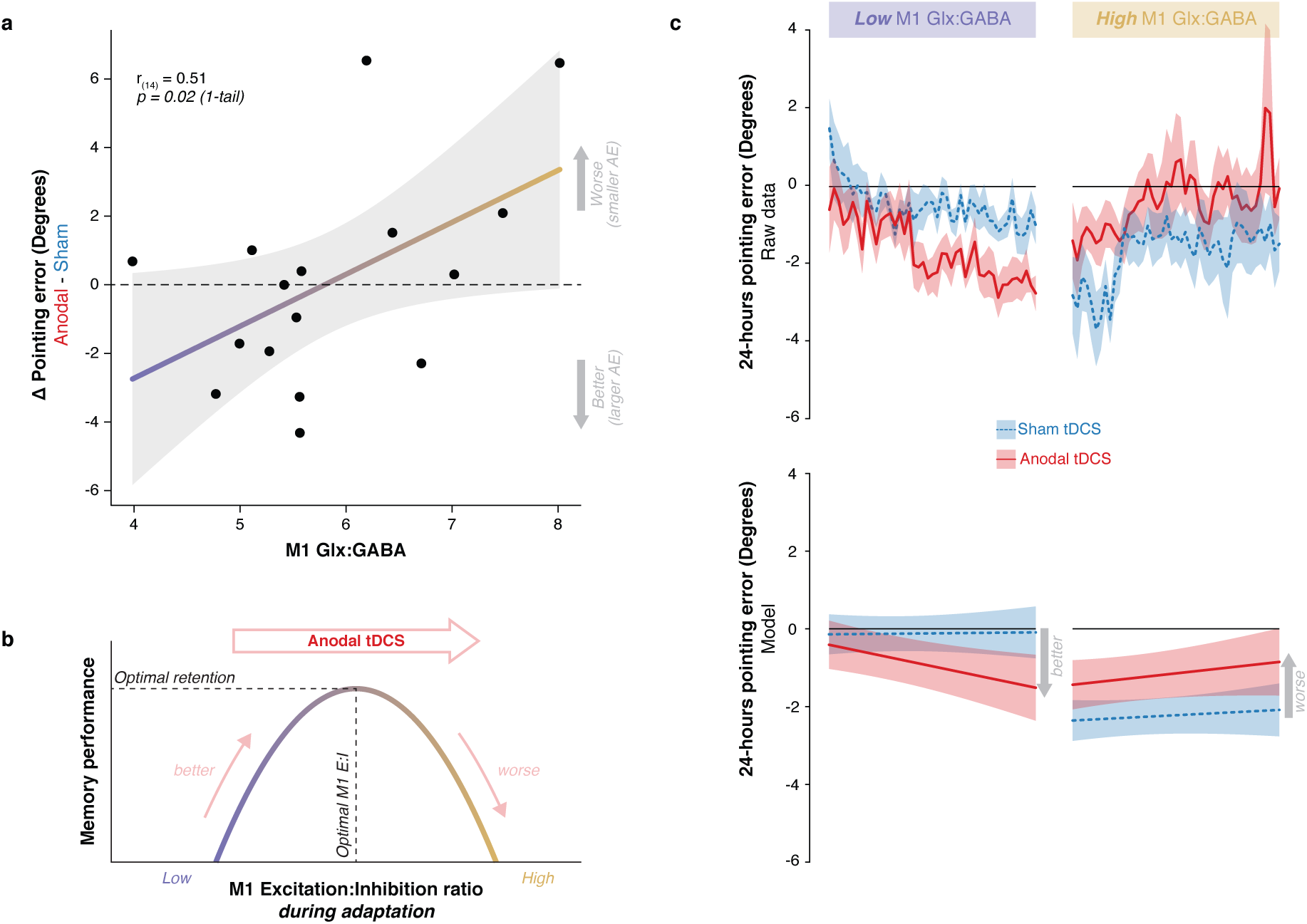
How stimulation changes memory depends on motor cortical E:I. **a.** Individuals’ M1 E:I (Glx:GABA) is plotted against the stimulation effect (anodal - sham difference in normalised pointing error at 24-hour retention). On the y-axis, negative values indicate greater retention with anodal tDCS compared to sham. Positive values indicate the opposite. Across individuals, stimulation enhanced retention in those with low E:I and impaired retention in those with high E:I. These data confirm the hypothesis that retention depends causally on M1 E:I. **b.** The schematic offers an explanatory account of the data in panel a. Under the assumption that stimulation increases E:I across the group, the distribution of induced memory change has an inverted U-shape. This suggests there is an optimal range of E:I within which retention is maximal. The optimum differs across individuals. By increasing E:I, stimulation moves individuals with low E:I towards maximum, increasing retention. But for individuals with high E:I, who are close to maximum, stimulation exceeds the optimum and so retention becomes impaired. **c.** A moderation analysis confirmed that how stimulation changed memory varied as a function of M1 E:I (Glx:GABA × tDCS : *t*_1419_ = 2.40, *p* = 0.017). For visualisation purposes, this interaction is illustrated using a median split on the M1 E:I data. The data (top) and model fit (bottom) are plotted separately for individuals with low versus high M1 E:I, and show opposing effects of excitatory stimulation on adaption memory that depend on individuals’ M1 E:I

Figure 6 plots the result of the moderation analysis for long-term retention. Panel a shows how the induced memory change varied as a function of M1 E:I – in those individuals with low E:I, stimulation enhanced retention; in individuals with high E:I, stimulation impaired retention. Panel b offers an explanatory account. Under the assumption that M1 a-tDCS increases E:I^50–52, 58^, the pattern of induced memory change followed an inverted U-shaped distribution, which suggests there is an optimum level of E:I at which retention is maximal. Increasing E:I via stimulation enhanced memory in those with low E:I, up to an optimum level beyond which stimulation had a deleterious effect, impairing retention. Panels c & d illustrate the result of the moderation analysis via a median split on the M1 E:I data.

A follow-up LMM decomposed the E:I data to assess the moderating roles of M1 GABA and Glx separately. Both Glx (Glx × a-tDCS: *t*_(1415)_ = 2.57, *p* = 0.005, one-tail, 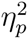 = 0.29) and GABA (GABA × a-tDCS: *t*_(1415)_ = −1.73, *p* = 0.042, one-tail, 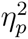 = 0.16) moderated the stimulation effect, each in opposite directions (Table S9 - model 6). Across individuals, stimulation increased retention in those with higher GABA and/or lower Glx, and impaired retention in those with lower GABA and/or higher Glx. This result was unchanged when controlling for average movement speed during prism exposure (Table S10). It was also anatomically specific: V1 neurochemistry did not moderate the effect of stimulation on retention (Table S11).

Given the finding in Experiment 1 of an association between age and M1 E:I (Fig. 2), we tested whether age could be substituted as a simpler, easy to measure, proxy for neurochemistry. By contrast with M1 E:I, age alone did not moderate the effect of stimulation on retention (see Supplementary Results; Fig. S3).

### How stimulation changes memory did not interact with genotype

Previous work in young adults showed that the enhancing effect of M1 a-tDCS on motor skill memory was influenced by a common polymorphism in the gene that codes for brain derived neurotrophic factor (BDNF val66met ^59^). Here we tested for a similar effect on adaptation memory. BDNF polymorphism (obtained in 24 participants in Experiment 2) did not significantly moderate the effect of M1 a-tDCS on retention (see Supplementary Results; Fig. S4).

## Discussion

This study tested the hypothesis that healthy older adults would exhibit stronger adaptation memory, owing to age-related M1 GABA decline. The results confirmed this prediction. In a cross-sectional study of healthy men (aged 49-81), older adults had higher long-term retention (Fig. 1) and lower M1 GABA (Fig. 2). A mediation analysis showed that the latter explained the former (Figs. 3 & 4). When M1 neurochemistry was accounted for, there was no longer a relationship between age and memory, consistent with full mediation. The findings were specific: anatomically (M1 not V1), neurochemically (GABA not Glx) and functionally (long-term not short-term retention). To more directly infer a causal link between neurochemistry and memory, anodal tDCS was used to experimentally lower M1 inhibitory tone^50, 52, 58^ (thus increasing E:I) in a subset of the same participants. Individuals’ intrinsic excitation:inhibition ratio within sensorimotor cortex moderated how stimulation changed memory (Fig. 6). In those with naturally low E:I (low Glx and/or high GABA) stimulation increased retention, whereas in those with naturally high E:I (high Glx and/or low GABA) stimulation impaired retention. The distribution of stimulation-induced memory change fitted an inverted U-shape, suggesting there is an optimum range of M1 E:I within which adaptation memory is maximal. The results were specific to E:I within M1 (no effect for V1). Whereas GABA loss in older age has typically been associated with functional decline^23, 26, 31, 32^, the present results reveal a specific domain of motor function that instead becomes naturally enhanced. Enhanced adaptation memory may help compensate for impaired motor skill learning in older age^60^.

This study extends our previous behavioural findings that M1 a-tDCS during adaptation enhances retention, thus boosting therapeutic efficacy in post-stroke visual neglect^48^. The present work provides mechanistic insight into the neurochemical basis of the induced memory gain. The results indicate that adaptation memory strength, both intrinsic and experimentally induced, depends causally and inversely on M1 cortical inhibitory tone. Previously we provided preliminary evidence for this hypothesis from an experiment in a small sample (*n* = 10) of young healthy adults. We inferred a likely causal link between M1 GABA and memory strength because stimulation-induced changes in both were correlated. Here, in a larger sample of older adults (a more relevant age for stroke translation) we provide more robust causal evidence for this hypothesis from mediation and moderation analyses of correlational and interventional data.

Two manipulations, one natural and one experimental, indicate that adaptation memory depends causally on M1 E:I - ageing (Fig. 4) and brain stimulation (Fig. 6). Collectively, they show that M1 E:I determines retention. On average, lower inhibitory tone is associated with stronger retention. However, the inverted U-shaped stimulation response (Fig. 6) suggests that there is an optimal level of E:I at which retention is maximal. This optimum differs across individuals. Increasing E:I via stimulation moves individuals with naturally low E:I towards their maximum, whereas individuals with naturally high E:I exceed that maximum and retention becomes impaired. On average, relatively younger (old) adults are more likely to have E:I levels below this theoretical upper physiological bound, while older (old) adults are closer to it. This may explain the absence of a significant overall group mean memory enhancement effect of stimulation in the present sample of older adults (Fig. 5), by contrast with our previous findings in young adults^48^. This suggests that, in general, inducing plasticity with M1 a-tDCS is, on average, likely to be less effective in older adults – at least to the extent to which it depends on lowering inhibitory tone^52, 61^. Nonetheless, age *per se* did not predict response to stimulation (Fig. S3), whereas M1 E:I did, underlining the potential utility of M1 Glx/GABA as biomarkers of inter-individual variation in stimulation response. The present results reveal that adaptation memory and M1 a-tDCS effects share a common neurochemical substrate: causal dependence on M1 inhibitory tone. This mechanistic synergy makes M1 a-tDCS a particularly suitable manipulation for understanding retention of adaptation, and vice versa.

Rather than enhance retention, stimulation impaired it in individuals with intrinsically high M1 E:I (Fig. 6a). An alternative interpretation of this result (to that offered above, Fig. 6b) is that M1 a-tDCS may have *reduced*, rather than increased, E:I in those individuals, contrary to the effect predominantly reported in the literature^50–52, 58^. Our prior work indicates that this is possible^48^. If, in those individuals with high M1 E:I, excitation is already near physiological ceiling, then anodal stimulation could trigger homeostatic regulatory mechanisms (that protect against over-excitation) to instead reduce E:I^62–66^ (Fig. S5). Thus, impaired retention would not be explained by the falling part of the inverted U-shaped [retention × M1 E:I] function that we propose in Fig. 6b. Rather, it would arise from E:I rebound via homeostasis when stimulation causes the excitability ceiling to become breached (Fig. S5). The relative (non-exclusive) contributions of these two potential mechanisms to impaired retention is a question for future work. Nonetheless, under either scenario, the data provide causal evidence that there is an optimal range of M1 E:I, that varies across individuals, within which retention is maximal.

In our previous work on neglect^48^, we speculated that M1 a-tDCS might antagonize (rather than enhance) prism adaptation therapy in some patients. The present data support this since here stimulation disrupted retention in healthy older individuals with naturally higher M1 E:I. Stroke disrupts E:I across distributed cortical networks, and how this interacts with age-related dysregulation is likely to vary by region and time. How the neurochemical constraints identified here apply in stroke populations remains to be tested. The present normative dataset could help guide interpretation of future stroke data. Serendipitously, in our earlier work, we established proof of concept via experiments in young adults^48^. These showed that M1 a-tDCS during adaptation specifically enhanced retention. If instead we had started by testing older healthy controls, we are unlikely to have ever progressed to testing prism therapy + M1 a-tDCS in neglect, since (as in Fig. 5) we would not have found evidence that stimulation enhances retention. What reconciles our previous and present results, across the combined evidence from younger and older adults, is the new physiological insight that individuals motor cortical E:I – both intrinsic and induced – governs this form of motor memory plasticity.

While the results provide a conceptual replication and extension of our previous finding that intrinsic differences in neurochemistry explain inter-individual variability in stimulation response, a critical caveat is that we sampled only men. This choice was informed by the fact that GABA levels change across the menstrual cycle in women and hence adaptation memory, E:I, and stimulation responsivity are also likely to fluctuate accordingly. Given hormonal changes across the lifespan, womens M1 inhibitory tone may have a different or more variable age-related trajectory than men. Hence, to rule out these hormonal sources of variance, which would require much larger samples, we did not recruit women. In so doing, we follow a long tradition in biomedical research, the limitations and adverse consequences of which for women are significant^67^. Of note, in our previous work, all three patient cases (by coincidence) were men. This may matter for the therapeutic effects observed. By excluding gender-related heterogeneity, we could identify an important mediator (M1 E:I) of variation in response to stimulation-induced functional plasticity, at least in men. Dissecting out intrinsic biological factors in this way helps to causally explain inter-individual differences and to dispel scepticism that this variability somehow renders brain stimulation (and tDCS in particular) suspect as a neuroscience tool^68^. We also assessed genotypic variation. Unlike previous work on motor skill retention^59^, we did not find that the BDNF val66met genetic polymorphism influenced the effect of M1 a-tDCS on retention of adaptation, at least not in this sample of older adult men (Fig. S4).

Our previous behavioural findings causally implicated sensorimotor cortex in retention because, on average, M1 a-tDCS during adaptation led to stronger memories. Yet the spatially diffuse electric field induced by tDCS complicates functional localization. The current work provides complementary evidence at the level of individuals that differences in adaptation memory relate to differences in M1 neurochemistry. This need not be interpreted as evidence that memories are formed and/or stored locally and/or exclusively in sensorimotor cortex. Of note, we measured brain chemistry only in M1 and V1. Adaptation memory, like most functions, is likely to be distributed,implemented through parieto-premotor-cerebellar circuit interactions. Yet we targeted M1 owing to evidence that it has a causal role in the early consolidation of motor learning^69–73^. These data strengthen this evidence base in the case of adaptation. We interpret the data to indicate that M1 is a privileged node in the distributed cortical circuitry that implements the early formation of adaptation memory. That is, the strength of that memory trace can be changed during its formation by tonic disinhibition of M1 (via a-tDCS), and the impact on individuals memory is quantitatively and causally related to their local E:I balance within M1. This local neurochemical measure has been shown to correlate with sensorimotor network resting state functional connectivity^74, 75^. Thus, M1 E:I may be an informative measure because it also serves as a proxy readout of sensorimotor network strength. Extra-synaptic GABAergic tone, that measured by magnetic resonance spectroscopy, has also been linked to oscillatory markers of inter-regional neuronal communication^76, 77^. Hence, this local M1 readout may also indirectly index inter-individual differences in propensity for inter-areal communication strength, of functional relevance during adaptation. Thus, we conclude that M1 is a sensitive node at which to both measure and manipulate adaptation memory formation.

Previous work has generally found adaptation to be preserved or somewhat impaired in older adults^35–46^. Instead, our work reveals that a specific sub-domain of adaptation – long-term memory – is naturally enhanced in older adults and provides a neurochemical explanation of this phenomenon. Whether these findings are specific to reach adaptation, or prisms, or may also generalize to other effectors and forms of adaptation remains to be investigated. Regardless, they are relevant for translational research on ageing and stroke rehabilitation. Adaptation has been regarded as of limited value for rehabilitation because memory for what is learned decays so quickly^78^. Hence, much research effort (including by us) has been invested in developing neuroscience interventions to boost retention^47, 48, 79–81^. Our finding that retention is naturally upregulated in (healthy) older age suggests such effort might be misplaced. If retention is already boosted naturally by ageing, then optimizing training regimens to exploit this for functional gain may instead prove a more profitable focus. This applies to health and disease. For example, postural imbalance, a cause of falls in older age, is associated with M1 GABA decline^31, 32^. Balance board training, used to counteract this, might be enhanced by incorporating an adaptation component, with the logic that the same GABA decline could promote retention of adapted training effects. Similarly, random as opposed to blocked practice impairs motor learning but boosts retention^82, 83^. Leveraging this psychological insight, while capitalizing on naturally greater retention of adaptation in older age, may inspire the design of novel training regimes that better promote the maintenance of motor skills and thus functional independence through older age.

These results identify a domain of upper limb motor functional plasticity that is naturally increased in healthy ageing. Is this likely to be beneficial? That will vary with context. Adaptation adjusts behaviour to counteract perturbations that impair performance, thus maintaining motor success. Once adapted, the optimal timescale for retention is the one that matches that of the perturbation^84^. For long-lasting, slowly evolving perturbations, such as gradual muscle stiffening with increasing age, adapting to that and maintaining it over time would help to offset these deleterious effects. Conversely, in volatile environments that require agents to quickly learn and forget new visuomotor transformations (e.g., playing basketball on a windy day), slow forgetting would be maladaptive. Hence, whether reduced M1 inhibitory tone and stronger retention is adaptive or maladaptive depends on the context and the task. Higher GABAergic tone (in physiologically younger adults) may enable more selective functional release of inhibition during learning^30^ and thus promote more selective retention (e.g. of those motor memories likely to be useful in future). Hence, stronger adaptation memory in older age is best conceived of as a “paradoxical functional facilitation”^85^ – an isolated domain of upregulated function that may have benefits, but which is a side effect of a deleterious process (age-related GABA loss) that primarily causes functional decline^23, 26, 31, 32^

Nonetheless, our findings provide grounds for optimism about healthy motor ageing. The usual narrative is one of decline and loss. Here instead we identify a natural age-related functional gain. Maybe we cannot “*teach an old dog new tricks*”, but we can instead focus effort on adapting and retaining existing skills, promoted by natural neurochemical changes that may contribute to maintaining motor function for longer.

## Materials and Methods

### Participants

Thirty two right handed men aged between 49 and 81 (mean age: 67.5 years, *s.d.*: 8.1) participated in this study. All were screened to rule out any personal or family history of neurological or psychiatric disorder and safety contraindications for the MRS and tDCS measurements. Written informed consent was provided by all participants. The study was approved by the U.K. NHS Research Ethics Committee (Oxford A; REC reference number: 13/SC/0163). In Experiment 1 all participants (*n* = 32) performed prism adaptation and tests of short-(10-minutes) and long-term (24-hours) retention. A sub-sample underwent a magnetic resonance spectroscopy scan to measure neurochemistry in left sensorimotor cortex (*n* = 22) and in an anatomical control volume in occipital cortex (*n* = 20; Fig. S2). A sub-sample consented to also participate in Experiment 2 (*n* = 25), consisting of two weekly sessions of prism adaptation combined with anodal/sham tDCS to M1. Full details of which measurements were obtained for each individual are in Table S1.

In Experiment 1, the sample size (n = 32) was chosen for comparability with previous MRS studies that successfully identified associations between behaviour and age-related GABA change^23, 26, 28, 29, 86^. Sample sizes in these previous studies ranged from 11 to 37. In Experiment 2, sample size was determined based on a power analysis run in G*Power^87^ (Version 3.1.9.2), informed by the stimulation effect size reported in our previous work^48^. In that study, left M1 a-tDCS enhanced long-term retention up to four days after adaptation, with an effect size of *d* = 0.73. The minimum sample size required to detect an effect of *d* = 0.73 with *α* = 0.05, and *β* = 0.95 was *n* = 22 (based on a one-tailed difference of two dependent means). To allow for potential dropouts, twenty-six participants were recruited. One participant was lost to retention follow-up and was therefore not included in the final sample of n = 25.

### Prism adaptation protocol

In both experiments, PA was performed using a purpose-built automated apparatus (Fig. S1a). Participants sat with their head fixed in a chinrest, viewing a 32-inch horizontal touchscreen through a liquid crystal display shutter. The touchscreen was used to present the visual targets and record reach endpoints. The shutter was used during after effect trials, and turned opaque at reach onset, thus occluding visual feedback of endpoint accuracy. The task was programmed in MATLAB version 2014b (MathWorks; https://uk.mathworks.com) using Psychtoolbox^88^ version 3, run on a MacBook Pro laptop. On each trial an audio voice recording instructed participants to reach and point with their right index finger at one of three targets on the touchscreen. There were two lateral targets situated 10 cm to the left and right of the central target. The distance between participants’ eyes and the central target was 57 cm.

During PA participants alternated between two types of task block: closed-loop pointing (CLP) and open-loop pointing (OLP). On closed-loop trials, participants wore 10 degree right-shifting prism goggles and were instructed to make rapid reaching movements (mean duration: 452 ms, *s.d.*: 119 ms) to either the left or right target in a pseudo-randomised order. Both speed and accuracy were emphasized. To limit strategic adjustments and “in-flight” error correction ^89, 90^ visual feedback of the first third of each reaching movement was occluded, as in previous work^48, 56, 91^. On open-loop trials, prisms were removed and participants were instructed to point at the central target (mean duration: 799 ms, s.d.: 135 ms). Accuracy was emphasized over speed. Visual feedback was prevented on each trial by the LCD shutter turning opaque at reach onset, thus occluding vision of the target, reach and endpoint error. This enabled the leftward after-effect to be measured without participants de-adapting in response to visual error feedback.

In both experiments, each PA session measured pointing accuracy during: baseline, adaptation, short-term (10-minutes) and long-term retention (24-hours; Fig. S1). Baseline closed- and open-loop pointing accuracy was measured in two blocks of 20 and 30 trials respectively. Adaptation comprised of alternating pairs of closed- and open-loop pointing blocks, six in Experiment 1 and seven in Experiment 2 (Fig. S1). Retention of the after-effect was measured 10-minutes and 24-hours after the end of PA, by means of a single block of 45 open-loop trials. In Experiment 2, 10-minute retention was followed by a washout phase in which participants pointed without wearing prisms, observed their leftward errors and therefore deadapted. Washout consisted of 40 closed-loop trials and 45 open-loop trials distributed across six interleaved blocks (Fig. S1b). The purpose of washout was to maximise the likelihood of observing a stimulation-related retention benefit at 24-hours. We reasoned that, if memory formation was strengthened by stimulation during adaptation, then washout should negatively interfere with long-term retention in the sham condition, but not in the anodal condition.

### Transcranial direct current stimulation

In Experiment 2, tDCS was delivered by a battery driven DC stimulator (Neuroconn GmbH, Ilmenau, Germany) connected to two 7 × 5 cm sponge electrodes soaked in a 0.9% saline solution. The anodal electrode was centred over C3 (5 cm lateral to Cz) corresponding to the left primary motor cortex according to the international 10-20 electrode System^92^. The cathode was placed over the right supraorbital ridge. During anodal tDCS, stimulation was applied at 1mA for 20 minutes, throughout the entire adaptation phase, as in our previous work^48^. The current ramped up and down over a 10 second period at stimulation onset and offset. During sham tDCS, the procedure was identical except that no stimulation was delivered during the 20 minutes. Instead, small current pulses (110 *µ*A over 15 ms) occurred every 550 ms to simulate the transient tingling sensations associated with real stimulation. Both experimenters and participants were blinded to the stimulation condition (anodal or sham) during behavioural testing. This was achieved by using blinding codes (“study mode” of the stimulator) provided by a researcher who was not involved in behavioural testing. Unblinding occurred at the statistical analysis stage, once data collection was completed. In Experiment 2, participants performed two PA+tDCS sessions (anodal/sham, order counter-balanced), each separated by a minimum of one week.

### MRS acquisition protocol

MRS data were acquired at the Oxford Centre for Clinical Magnetic Resonance Research (OCMR, University of Oxford), on a Siemens Trio 3-Tesla whole-body MR scanner and using a 32-channel coil. High resolution T1-weighted structural MR images (MPRAGE; 224 × 1 mm axial slices; TR/TE = 3000/4.71 ms; flip angle = 8°; FOV = 256; voxel size = 1 mm isotropic; scan time = 528 secs) were acquired for magnetic resonance spectroscopy (MRS) voxel placement and registration purposes. MRS data were acquired from two volumes of interest (VOIs) in two consecutive acquisitions. The first VOI was centred on the left motor hand knob^93^ and included parts of the pre- and post-central gyrus (Fig. S2c). The second (anatomical control) VOI was centred bilaterally on the calcarine sulcus in the occipital lobe (visual cortex)^94–96^ (Fig. S2c). B0 shimming was performed using a GRESHIM (64 × 4.2 mm axial slices, TR = 862.56 ms, TE1/2 = 4.80/9.60 ms, flip angle = 12°, FOV = 400, scan duration = 63 secs). MR spectroscopy data (spectra) were acquired using a semi-adiabatic localization by adiabatic selective refocusing (semi-LASER) sequence (TR/TE = 4000/28 ms, 64 scan averages, scan time = 264 secs) with variable power radio frequency pulses with optimized relaxation delays (VAPOR), water suppression, and outer volume saturation ^97, 98^. In addition, unsuppressed water spectra were acquired from the same VOIs to remove residual eddy current effects, and to reconstruct the phased array spectra ^99^. Single-shot acquisitions were saved separately (single-shot acquisition mode), then frequency and phase corrected before averaging over 64 scans.

### MRS data analysis

Metabolites were quantified using LCModel^100–102^ performed on all spectra within the chemical shift range 0.5 to 4.2 ppm. The model spectra were generated based on previously reported chemical shifts and coupling constants by VeSPA Project (Versatile Stimulation, Pulses and Analysis). The unsuppressed water signal acquired from the volume of interest was used to remove eddy current effects and to reconstruct the phased array spectra^99^. Single scan spectra were corrected for frequency and phase variations induced by subject motion before summation. Glutamix (Glx) was used in the current study due to the inability to distinguish between glutamate and glutamine using a 3T MRI scanner. To avoid biasing the sample towards high concentration estimates, an expected relative Cramér-Rao Lower Bound (CRLB) was computed for each individual dataset given the concentration estimate and assuming a constant level of noise across all measurements (see *SI* for detailed methods). Datasets for which the Pearson residual between the expected and observed relative CRLB exceeded 2 were excluded from subsequent analysis. Using this quality filtering criterion for *γ*-Aminobutyric acid (labelled GABA), Glutamix (Glutamine+Gutamate, labelled Glx) and total Creatine (Creatine + Phosphocreatine, labelled tCr), four V1 MRS datasets were discarded and no M1 MRS dataset was discarded.

Tissue correction is an important step in MRS data analysis, especially in older adults owing to brain atrophy^103^. LCmodel outputs metabolite concentrations for an entire volume of interest. So if the fraction of neural tissue within a volume of interest is low, owing to age-related atrophy ^104^, metabolite concentration estimates will also necessarily be low. Several tissue correction techniques have been proposed to account for this potential confound, with currently no consensus in the literature^105, 106^. Most of these techniques make assumptions about the distribution of the metabolite of interest within the different tissue compartments. However, such assumptions may not hold across the lifespan, as the normal ageing process may affect some compartments more than others. Hence, all analyses reported in this paper used non-tissue corrected concentration estimates and instead included the percentage of grey matter (GM) and white matter (WM) in the MRS voxel as confounding variables of no interest (as in ^107^). Since this partial volume correction approach makes no assumption about the distribution of GABA and Glx within the different tissue types, it is particularly suitable for the present study (in which participants ranged in age from 49 to 81), and hence controls for atrophy while remaining agnostic about the differential impacts of ageing on tissue types. The percentages of grey matter, white matter, and cerebrospinal fluid present in the VOIs were calculated using FMRIBs automated segmentation tool^108^. They are reported together with MRS data quality metrics in Table S2.

Across individuals, the total creatine (tCr) concentration estimate was negatively correlated with age in the M1 voxel (*r*_(22)_ = −0.46*, p* = 0.04) although not in the V1 voxel (*r*_(18)_ = −0.06*, p* = 0.81; Fig. S2b). Owing to this confound with age, tCr could not be used as a valid internal reference for metabolite estimates. Hence, throughout this work, we used absolute concentration estimates for GABA and Glx, rather than expressing the data as ratios of tCr.

### Statistical analysis

Statistical analyses of behaviour were performed in R^109^. To control for interindividual differences in pre-adaptation pointing accuracy, across all trials endpoint error data were normalized by subtracting the average pointing error at baseline (across left/right targets for closed-loop blocks; middle target for open-loop blocks). Unless specified otherwise, all statistical tests were two-tailed. Analyses were performed using linear regression. Linear mixed-effects models (LMMs) were used for analyses with a longitudinal/repeated-measures component (e.g. adaptation, retention) by including intercepts and slopes as participant random effects. This approach has two advantages compared to repeated measures analyses of variance (ANOVAs): it allowed us to 1) also consider within-block behavioural dynamics, as opposed to only block average errors, and 2) dissociate random sources of inter-individual variability from meaningful ones. All model specifications are reported in Supplementary Tables. We compared LMM model parameters directly to establish neuroanatomical and neurochemical specificity. Model parameters were compared using a general linear hypothesis test using the *multcomp* package in R^110^. For visualisation purposes, Figs. 1b, 3 and 6b show block-averaged data as measures of retention, but the statistical analyses were run on individual trial data with random intercepts and slopes. Measures of effect size are reported for all substantial analyses, using the *effectsize* package^111^ in R. Cohen’s *d* was used to compute effect sizes for a one-sample t-test against zero for short- and long-term retention in Experiment 1, and for paired-samples t-tests of sham versus anodal stimulation on short- and long-term retention in Experiment 2. Approximate partial eta-squared (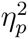) for linear mixed-effects regression analyses to summarise the proportion of variance associated with a particular fixed effect. Rules of thumb have been proposed for interpreting effect sizes. The norms for Cohen’s *d* are: small = 0.2; medium = 0.5; large = 0.8. The norms for 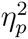 are: small = 0.01; medium = 0.06; large = 0.14^112^.

Because GABA is synthesised from glutamate, the concentrations of these two neurotransmitters are typically correlated positively in the brain^113, 114^ (in our dataset, M1 GABA × M1 Glx: *r*_(20)_ = 0.34; V1 GABA × V1 Glx: *r*_(14)_ = 0.16). Therefore, when analysing the relationship between the absolute concentration in GABA or Glx within a voxel and outcome, the concentration of the other neurotransmitter (GABA or Glx) was also included in the model. In addition, grey and white matter concentrations were also included as covariates of no interest in all models that included neurochemical data.

A mediation analysis was used to characterise the “mechanistic” links underlying the observed correlations between age, neurochemistry, and retention. This was performed using the R package *mediation* for causal mediation analysis^115^. Mediation was conducted using regression with nonparametric bootstrapping (10,000 resamples) to ascertain whether M1 inhibitory tone accounted for the link between age and long-term retention. The model included: age as the independent variable (X); absolute concentrations of M1 GABA and Glx as mediators (M_1_, M_2_); block-averaged retention at 24-hours as the dependent variable (Y) (block mean error normalised by the baseline for each individual), and controlled for the fraction of GM and WM in the M1 voxel (C_1_, C_2_). The percentage mediation (*P_M_*) was calculated as the fraction of total effect (c) accounted by indirect effects (ab_1_ or ab_2_).

## Data and Code Availability

All analysis code and data related to this paper are available on the Open Science Framework (https://osf.io/stkv2/).

## Acknowledgements

This study was supported by the NIHR Oxford Health Biomedical Research Centre. The views expressed are those of the authors and not necessarily those of the NHS, the NIHR or the Department of Health. PP was funded by a scholarship from the Marie Sklodowska-Curie Initial Training Network (Adaptive Brain Computations). GS was funded by an Australian National Health and Medical Research Council (NHMRC APP1104692) Early Career Fellowship. HJB is supported by a Principal Fellowship from the Wellcome Trust (110027/Z/15/Z). JOS is supported by a Sir Henry Dale Fellowship from the Royal Society and the Wellcome Trust (HQR01720). The Wellcome Centre for Integrative Neuroimaging is supported by core funding from the Wellcome Trust (203139/Z/16/Z). For the purpose of Open Access, the authors have applied a CC BY public copyright licence to any Author Accepted Manuscript version arising from this submission. We thank Rebecca Annells for her help collecting data in Experiment 1.

## Supplementary Information

### Supplementary methods

#### MRS data quality filtering procedure

To treat noise in MRS data, ongoing methods development aims to develop procedures that systematically identify bad quality data in an objective way that does not depend on visual inspection of spectra by an expert. Over the years, the method of Cramér-Rao Lower Bounds (CRLB) has become the gold standard for determining concentration estimate uncertainty ^116^. The *CRLB* represents the lowest possible standard deviation of all unbiased concentration estimates obtained from fitting the model to the data. It is typically used in its *relative* form — labelled *%CRLB* — as a percentage of the estimated metabolite concentration. Its calculation for all estimated metabolites is part of the standard analysis pipeline of LCModel^100, 101^. Many authors have used the *%CRLB* as a way to perform quality filtering by rejecting metabolite concentration estimates for which the *%CRLB* falls above a certain arbitrary threshold (usually between 20% and 50%), which is judged as an unacceptable level of uncertainty in the estimate^?, 51, 58, 117, 118^.

Recently, however, some authors have warned against the usage of *%CRLB* for quality filtering because it could lead to wrong or missed statistical findings^119, 120^. For samples with large levels of noise, such as caused by bad quality MRS acquisition (e.g. too small voxel size, not enough averages, bad shimming) or bad quality spectrum fitting (e.g. inappropriate basis files), metabolite concentration estimates would be associated with high *%CRLB*, in a way that truly reflects high estimation uncertainty. In this scenario, it would be valid to mistrust the data based on a high *%CRLB*. However, because of the *relative* nature of the *%CRLB*, this metric also strongly depends on its denominator, i.e. the estimated metabolite concentration. Hence, for two samples acquired with equivalent levels of noise, the sample with a lower metabolite concentration will have a higher *%CRLB*. In that scenario, rejecting the dataset based on interpreting its high *%CRLB* as an indicator of high estimation uncertainty would be invalid.

This is relevant to the present study, in which we aimed to measure a reduction in GABA concentration with older age. The *%CRLB* cutoff criterion introduces a potential selection bias that could artificially bias the sample towards excluding participants with low GABA concentrations (and correspondingly high *%CRLB*). Therefore, to avoid this potential methodological confound we developed a data quality filtering approach that did not solely consider high *%CRLB* with respect to an arbitrary cutoff, but also considered the concentration estimate itself when deciding whether or not to reject datasets for quality control. Thus we aimed to better deal with the following scenarios: 1) datasets with a high *%CRLB* because of a low concentration estimate, rather than an excessive level of noise – such datasets should not be excluded; 2) Datasets with a low *%CRLB* simply because the concentration estimate is high might in fact be excessively high, given the metabolite concentration – such datasets should be excluded.

We therefore used the following method as an alternative to standard *%CRLB*-cutoff-based quality filtering. First, the following model was fitted to the “concentration estimate *× %CRLB*” relationship:

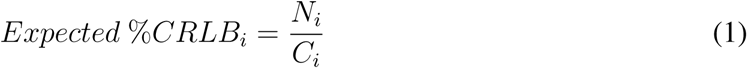

where *N_i_* represents a group noise constant and *C_i_* the concentration estimates for a metabolite *i*. Across the group, if this simple model can explain most of the variance in the observed relationship between concentration estimates and *%CRLB*, it means that the level of noise is relatively constant across all measurements. Any deviation from this model reflects an *unusual* level of noise compared to the other measurements. For each measurement, deviation from the model can be expressed as the Pearson residual as follows:

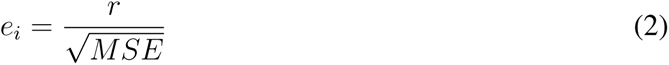

where *r_i_* is the raw residual (i.e. difference between the *%CRLB* and *expected %CRLB* for a certain measurement) and *MSE* is the mean squared error (i.e. mean deviation of all measurements from the model). The greater the Pearson residual for a given measurement, the noisier it is with respect to the rest of the data, irrespective of the concentration estimate. Note that this method does not reject the lower tail of the distribution entirely and therefore does not induce a selection bias towards high concentration estimates. Datasets with a Pearson residual greater than 2 were considered excessively noisy and were excluded from statistical analysis.

### Supplementary Results

#### Neurochemistry significantly moderates how stimulation changes retention but age does not

Given the inter-relationships between age, neurochemistry and retention we observed in Experiment 1 (Figs. 2-4),we also tested whether age (and not just M1 E:I) moderated how stimulation changed memory (in Experiment 2), and hence could serve as a (cheaper and easier to measure) surrogate of M1 E:I to predict stimulation response.

Age did not moderate the effect of stimulation on retention (age *×* a-tDCS: *t*_(1419)_ = 0.79, *p* = 0.43, two-tail). There was no association between age and the effect of a-tDCS on long-term retention (Fig. S3). This was the case when the full sample of all participants in Experiment 2 was analysed (*n* = 25; *r*_(23)_ = *−*0.12, *p* = 0.56). It was also true of the reduced sample of participants for whom we also had MRS data (*n* = 16; *r*_(14)_ = 0.19, *p* = 0.47). To directly compare the relative impacts of age versus M1 E:I on the behavioural effect of a-tDCS, we used Fisher’s r-to-z transform to contrast their Pearson correlation coefficients (i.e. relationships in Fig. 6a versus Fig S3). The relationship between the stimulation effect and M1 E:I was significantly greater than that between the stimulation effect and age for the full dataset (*t*_(14)_ = *−*2.53, *p* = 0.02). For the reduced sample, correlations did not differ significantly, presumably reflecting lower statistical power (*t*_(14)_ = *−*0.97, *p* = 0.35). We conclude that mere chronological age does not predict the effect of stimulation on retention, by contrast with M1 E:I.

#### Effect of BDNF polymorphism on the stimulation effect

Brain-derived Neurotrophic Factor (BDNF) is important for synaptic plasticity induction and has been shown to mediate the effect of anodal direct current stimulation^59, 121^. Individuals with the BDNF val66met polymorphism exhibit reduced behavioural and neural markers of motor cortical plasticity ^59, 122–124^. The polymorphism causes a partial reduction in activity-dependent BDNF secretion, and consequently reduces longterm potentiation^121, 125, 126^. Work in mice and humans has shown that plastic enhancement of motor skill retention via M1 anodal tDCS is reduced in val66met allele carriers ^59^. Hence, in this supplementary analysis, we acquired genotyping information to test whether BDNF polymorphism moderates the effect of M1 a-tDCS on retention of adaptation.

Genotyping was acquired for 24/25 participants in Experiment 2. Genomic DNA was extracted from buccal cells using the ChargeSwitch ® gDNA Buccal Cell Kit (ThermoFisher Scientific, UK) and samples were genotyped in duplicate by LGC Genomics (LGC Group, UK). Rs6265 was the only polymorphism examined.

The val66met polymorphism is present in about one third of the Caucasian population ^127^. In our sample, 5/24 participants (21% of our sample) carried the Met allele (4 Val-Met participants, 1 Met-Met participant). BDNF polymorphism (Met-allele carriers vs. non-carriers) had no significant influence on the effect of a-tDCS on either short-term (BDNF × tDCS × trial : *β* = −0.001, 95% CI [−0.014, 0.012], *t*_(2141)_ = −0.206, *p* = 0.84), Fig. S4) or long-term retention (BDNF × tDCS × trial : *β* = −0.01, 95% CI [−0.02, 0.002], *t*_(2141)_ = −1.61, *p* = 0.11), Fig. S4).

## Supplementary Figures

**Figure S1:**
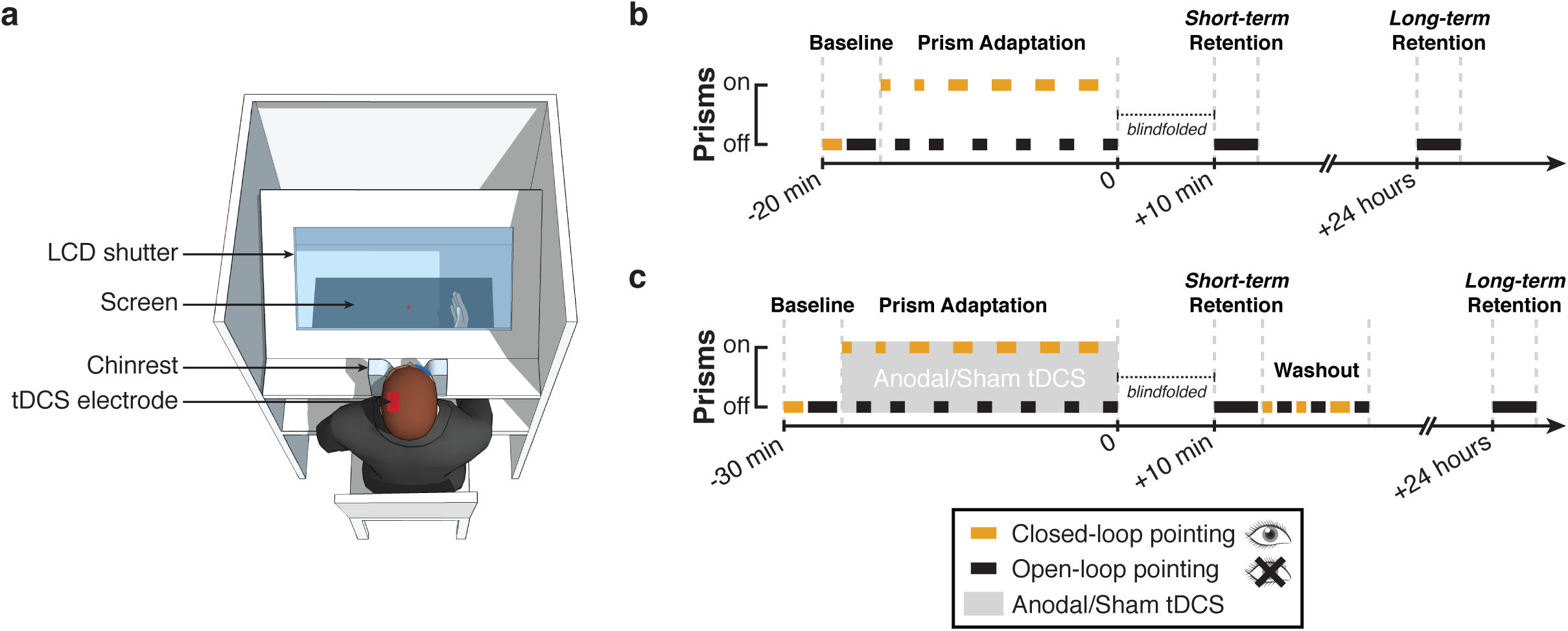
Prism adaptation protocol. a. Experimental setup. For both Experiment 1 and 2, participants sat in a chinrest viewing a horizontal 32-inch touchscreen through a liquid crystal shutter. The touchscreen was used to present visual targets and record reach endpoints. The liquid crystal display shutter was used to control visual feedback by turning opaque during reaching movements to conceal endpoint performance. **b. Procedure for Experiment 1.** Baseline accuracy was measured without prisms during blocks of closed-loop (continuous visual feedback) and open-loop (no visual feedback) pointing. During adaptation, participants alternated between blocks of prism exposure (closed-loop, glasses on) and after-effect measurement (open-loop, prisms off). Retention of the after-effect was measured 10 minutes and 24 hours post-adaptation. **c. Procedure for Experiment 2.** The procedure for Experiment 2 was the same as Experiment 1, except that left M1 anodal tDCS (real/sham) was applied throughout adaptation (grey shading). Short-term retention was followed by washout, during which participants observed and corrected their leftward errors (closed-loop pointing blocks, no prisms), interleaved with open-loop measures to confirm aftereffect decay back to baseline.

**Figure S2:**
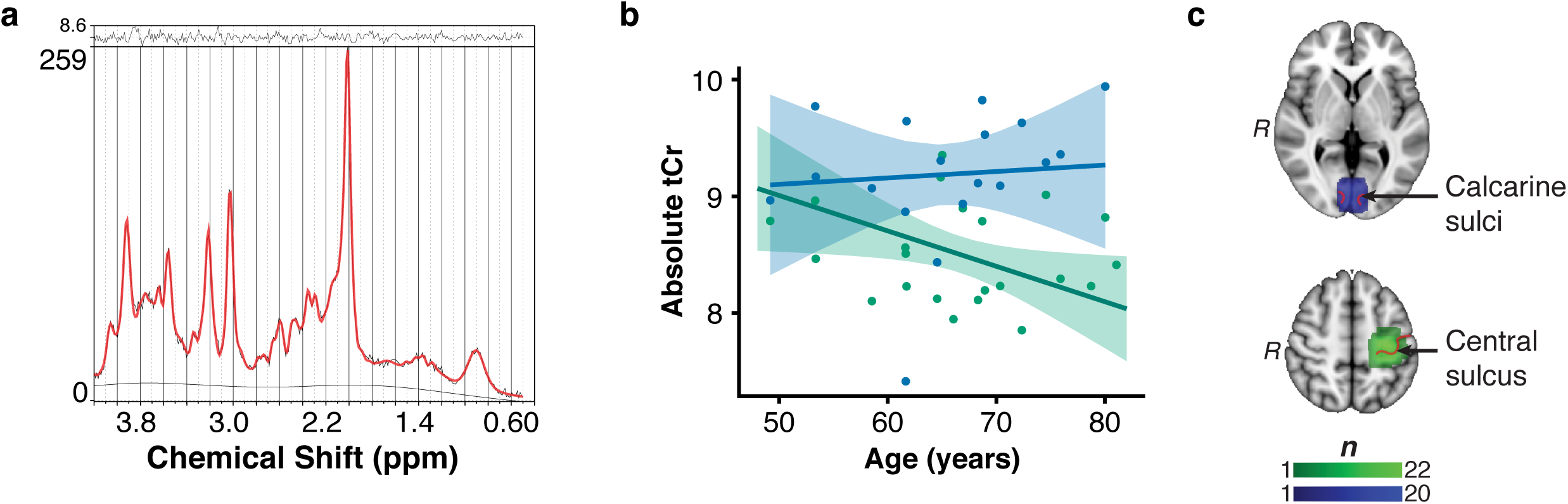
Magnetic resonance spectroscopy data quality. This figure shows the quality of MRS data collected in Experiment 1. **a.** Example raw MRS spectrum and LCModel fit from one participant. The fitted LCModel (in red) is plotted overlaid on the raw data (in black). The difference between the data and model (residuals) is shown at the top and the baseline is shown at the bottom. **b.** This panel presents the association between age and total Creatine (tCr), controlling for the fraction of WM and GM, in the M1 voxel (in green) and the V1 voxel (in blue). Shading indicates 95% confidence intervals. This panel shows that the tCr estimate was negatively correlated with age in M1, but not in V1. Because of this relationship, we use absolute concentrations of GABA and Glx throughout the paper, rather than using tCr for internal referencing. **c.** Magnetic resonance spectroscopy voxels group overlap map. The M1 voxel was centred on the left central sulcus in 22 participants (in green, MNI coordinate z = 52). The control V1 voxel was centred on the bilateral calcarine sulcus in 20 participants (in blue, MNI coordinate z = 2). Colour bar represents the degree of overlap. All images are displayed in radiological convention (i.e. left side of the image corresponds to the right side of the brain).

**Figure S3:**
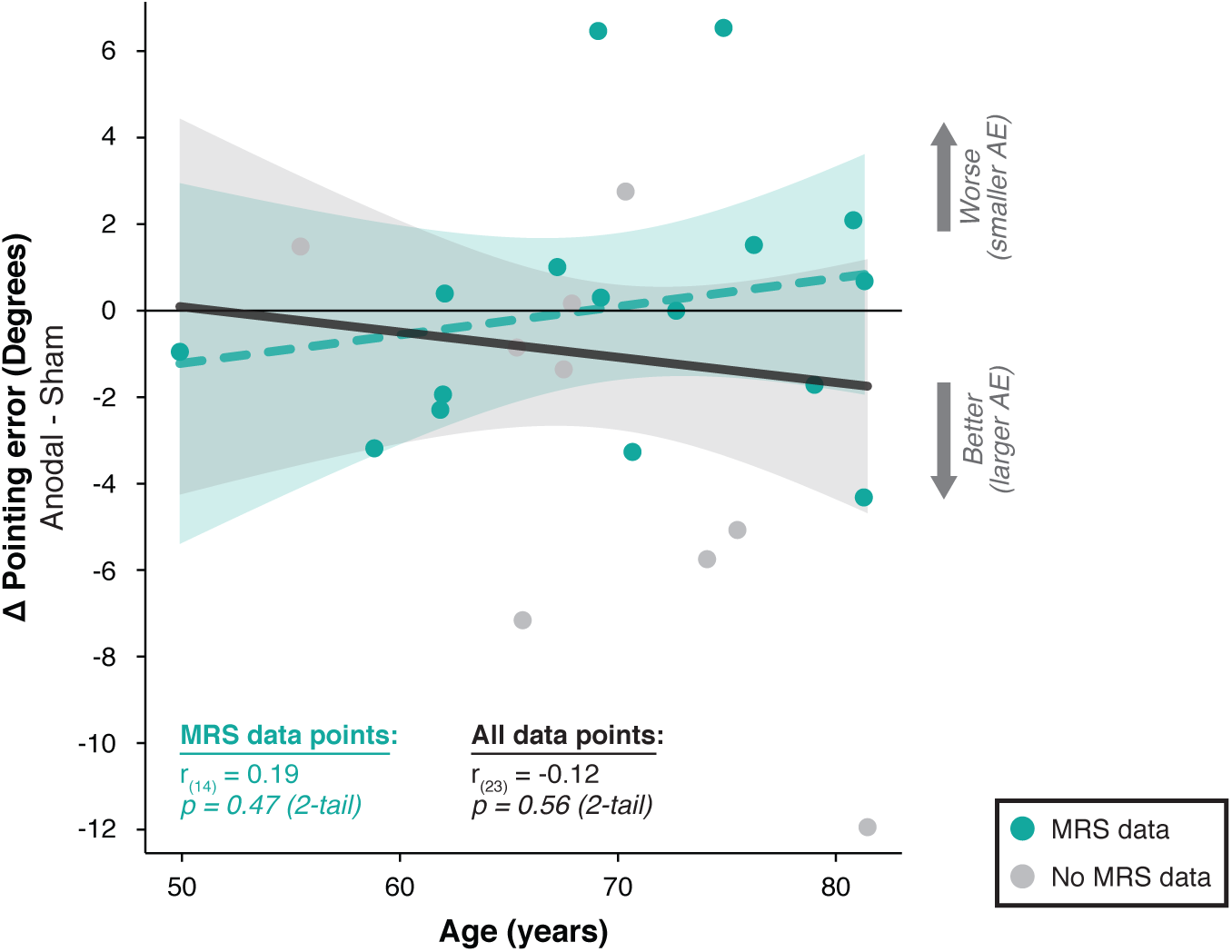
The behavioural effect of a-tDCS on long-term retention is not moderated by age. Age is plotted against the stimulation effect (anodal - sham difference in normalised pointing error at 24-hour retention). On the y-axis, negative values indicate greater retention with anodal tDCS compared to sham. Positive values indicate the opposite. The linear regression line is plotted for both the reduced sample of participants who had both MRS and retention data (*n* = 16, blue data points and line) as well as the full sample – including those participants who had missing MRS data (*n* = 25, black line). Irrespective of the sample considered, there was no association between age and the effect of a-tDCS on long-term retention (both *p >* 0.05).

**Figure S4:**
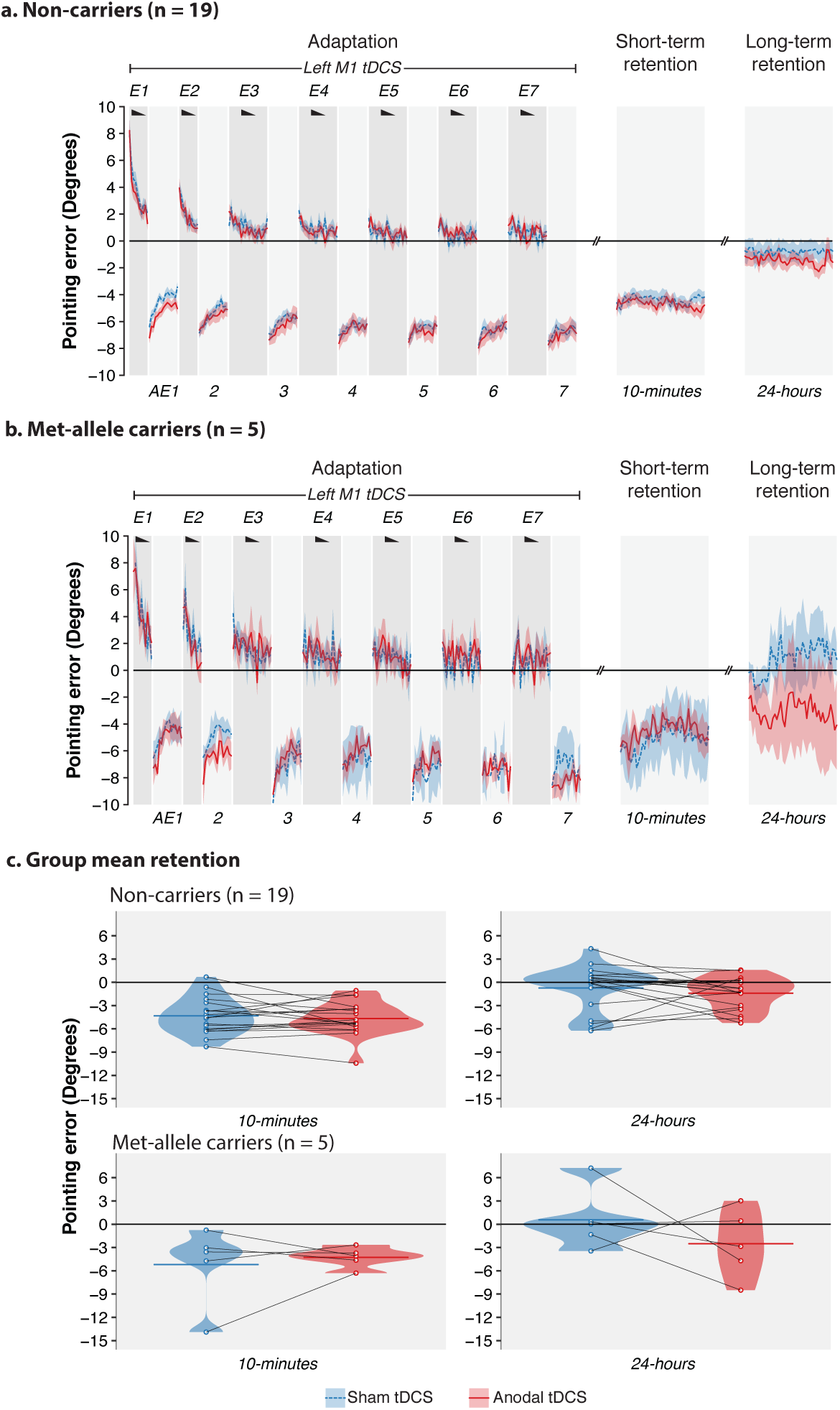
BDNF Val66Met genotype did not influence how stimulation changed retention. Plot shows the same data with the same conventions as in Fig. 5, separated by genotype. **a.** Behaviour of individuals with val/val genotype. **b** Behaviour of individuals with val/met genotype. **c.** Mean, distribution and individual datapoints for short-term (10-min) and long-term (24-hour) retention separated by genotype. Linear mixed models showed that BDNF val66met genotype did not significantly influence how stimulation changed short (BDNF × a-tDCS: *t*_(2141)_ = 1.00, *p* = 0.320) or long-term retention (BDNF × a-tDCS: *t*_(2141)_ = −1.61, *p* = 0.107).

**Figure S5:**
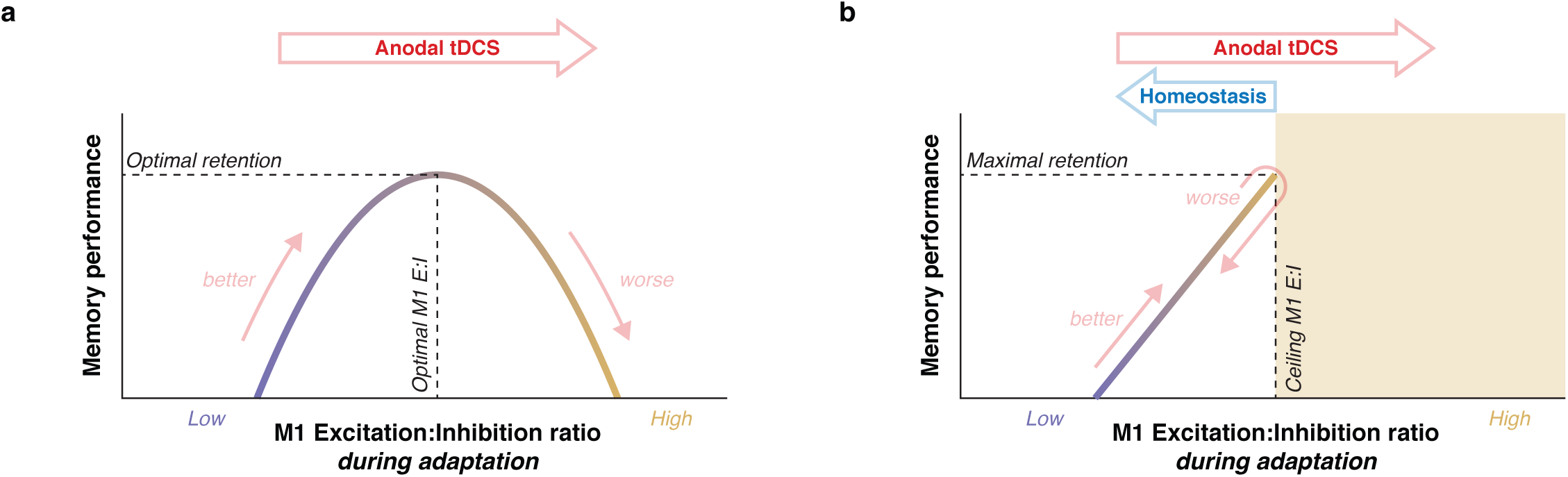
Alternative account of how stimulation impairs adaptation memory in individuals with high M1 E:I. a. For comparison purposes, the model proposed in Fig. 6b is reproduced here. **b.** The schematic offers an alternative mechanistic interpretation of the data presented in Fig. 6a. This model assumes that in a healthy brain there is a ceiling on cortical excitation which, when exceeded, triggers homeostatic mechanisms that reduce E:I, to bring it back within physiological range. In individuals with naturally high M1 E:I (near ceiling), homeostasis could overshoot, leading to an overall decrease in E:I. This mechanism could explain why stimulation impairs retention in those with high baseline M1 E:I, without requiring a reversal in the sign of the relationship between M1 E:I and retention.

## Supplementary Tables

**Table S1:** Participant demographics and inclusion details. Experiment 1 consisted of a behavioural experiment (prism adaptation, PA; with retention probe the next day) and a MR Spectroscopy scan. Experiment 2 consisted of two PA sessions plus retention probes 24 hours later, separated by one week. Tick marks indicate which measures were obtained for which participants. In the MRS column, parentheses indicate incomplete data due to technical difficulties (missing control voxel).

| Participant | Age | Experiment 1 |  | Experiment 2 |
| --- | --- | --- | --- | --- |
|  |  | *PA session* | *MRS* | *PA+tDCS sessions* |
| 1 | 81.1 | ✓ | (✓) | ✓ |
| 2 | 78.7 | ✓ | (✓) | ✓ |
| 3 | 80.0 | ✓ | ✓ | ✓ |
| 4 | 75.9 | ✓ | ✓ | ✓ |
| 5 | 74.6 | ✓ | ✓ | ✓ |
| 6 | 72.3 | ✓ | ✓ | ✓ |
| 7 | 68.7 | ✓ | ✓ | ✓ |
| 8 | 66.9 | ✓ | ✓ | ✓ |
| 9 | 64.6 | ✓ | ✓ |  |
| 10 | 53.3 | ✓ | ✓ |  |
| 11 | 68.9 | ✓ | ✓ | ✓ |
| 12 | 80.7 | ✓ |  | ✓ |
| 13 | 70.4 | ✓ | ✓ | ✓ |
| 14 | 66.0 | ✓ | ✓ |  |
| 15 | 61.7 | ✓ | ✓ | ✓ |
| 16 | 68.3 | ✓ | ✓ |  |
| 17 | 58.5 | ✓ | ✓ | ✓ |
| 18 | 53.4 | ✓ |  |  |
| 19 | 64.9 | ✓ | ✓ |  |
| 20 | 61.7 | ✓ | ✓ | ✓ |
| 21 | 49.2 | ✓ | ✓ |  |
| 22 | 61.6 | ✓ | ✓ | ✓ |
| 23 | 65.0 | ✓ | ✓ | ✓ |
| 24 | 65.6 | ✓ | ✓ | ✓ |
| 25 | 75.4 | ✓ |  | ✓ |
| 26 | 74.1 | ✓ |  | ✓ |
| 27 | 67.8 | ✓ |  | ✓ |
| 28 | 65.2 | ✓ |  | ✓ |
| 29 | 55.4 | ✓ |  | ✓ |
| 30 | 67.4 | ✓ |  | ✓ |
| 31 | 70.3 | ✓ |  | ✓ |
| 32 | 70.9 | ✓ |  | ✓ |

**Table S2:** MRS data quality metrics. GM: Fraction of Grey Matter; WM: Fraction of White Matter; CSF: Fraction of Cerebrospinal Fluid; SNR: Signal/Noise Ratio; CRLB: CramerRao Bounds; FWHM: Full-Width Half Maximum; *: datasets that failed quality filtering.

| Participant | Voxel | GM | WM | CSF | FWHM | SNR | CRLB (GABA) | CRLB (Glx) |
| --- | --- | --- | --- | --- | --- | --- | --- | --- |
| 1 | M1 | 19.79 | 56.58 | 23.63 | 0.036 | 35 | 24 | 6 |
| 2 | M1 | 19.84 | 66.32 | 13.84 | 0.048 | 31 | 21 | 5 |
| 3 | M1 | 19.50 | 62.99 | 17.51 | 0.024 | 35 | 34 | 7 |
| 4 | M1 | 20.68 | 56.34 | 22.99 | 0.048 | 39 | 23 | 4 |
| 5 | M1 | 24.10 | 46.12 | 29.79 | 0.036 | 31 | 23 | 4 |
| 6 | M1 | 29.63 | 55.30 | 15.07 | 0.036 | 35 | 24 | 6 |
| 7 | M1 | 26.99 | 58.13 | 14.88 | 0.036 | 41 | 31 | 5 |
| 8 | M1 | 25.85 | 58.52 | 15.63 | 0.048 | 36 | 19 | 4 |
| 9 | M1 | 24.89 | 63.06 | 12.05 | 0.048 | 37 | 24 | 4 |
| 10 | M1 | 30.13 | 60.06 | 9.81 | 0.036 | 35 | 25 | 6 |
| 11 | M1 | 27.02 | 64.41 | 8.57 | 0.036 | 43 | 26 | 5 |
| 13 | M1 | 23.74 | 61.53 | 14.73 | 0.048 | 34 | 22 | 5 |
| 14 | M1 | 28.64 | 53.80 | 17.56 | 0.036 | 42 | 19 | 5 |
| 15 | M1 | 24.63 | 61.86 | 13.51 | 0.036 | 38 | 21 | 5 |
| 16 | M1 | 25.15 | 59.97 | 14.88 | 0.048 | 32 | 29 | 6 |
| 17 | M1 | 24.25 | 69.98 | 5.77 | 0.048 | 35 | 23 | 6 |
| 19 | M1 | 29.71 | 54.34 | 15.96 | 0.036 | 36 | 20 | 6 |
| 20 | M1 | 23.26 | 57.15 | 19.59 | 0.036 | 33 | 20 | 6 |
| 21 | M1 | 20.37 | 68.78 | 10.85 | 0.036 | 36 | 27 | 7 |
| 22 | M1 | 30.82 | 51.41 | 17.77 | 0.036 | 35 | 20 | 5 |
| 23 | M1 | 29.36 | 55.34 | 15.30 | 0.048 | 39 | 26 | 5 |
| 24 | M1 | 22.72 | 59.81 | 17.47 | 0.048 | 39 | 18 | 5 |
| 1 | V1 | *Not acquired* |  |  |  |  |  |  |
| 2 | V1 | *Not acquired* |  |  |  |  |  |  |
| 3 | V1 | 48.51 | 29.48 | 22.01 | 0.036 | 39 | 25 | 5 |
| 4 | V1 | 52.00 | 20.39 | 27.61 | 0.048 | 38 | 30 | 4 |
| 5 | V1 | 47.26 | 30.44 | 22.22 | 0.036 | 44 | 22 | 4 |
| 6 | V1 | 50.06 | 24.07 | 25.87 | 0.048 | 38 | 22 | 4 |
| 7 | V1 | 47.90 | 30.28 | 21.80 | 0.036 | 41 | 19 | 5 |
| 8 | V1 | 44.54 | 37.33 | 18.13 | 0.048 | 43 | 21 | 5 |
| 9 | V1 | 46.64 | 34.90 | 18.47 | 0.036 | 44 | 26 | 4 |
| 10 | V1 | 53.85 | 28.87 | 17.27 | 0.048 | 39 | 21 | 4 |
| 11 | V1 | 54.21 | 31.50 | 14.30 | 0.048 | 46 | 18 | 4 |
| 13 | V1 | 43.67 | 39.36 | 16.98 | 0.036 | 47 | 21 | 4 |
| 14 | V1 | 50.81 | 32.97 | 16.22 | 0.048 | 17 | 120* | 14* |
| 15 | V1 | 51.95 | 34.06 | 13.99 | 0.048 | 26 | 35* | 8 |
| 16 | V1 | 40.91 | 33.66 | 25.43 | 0.048 | 34 | 26 | 5 |
| 17 | V1 | 56.66 | 28.67 | 14.67 | 0.036 | 41 | 17 | 4 |
| 19 | V1 | 55.35 | 29.95 | 14.70 | 0.059 | 32 | 30* | 6 |
| 20 | V1 | 47.20 | 36.26 | 16.54 | 0.036 | 40 | 23 | 5 |
| 21 | V1 | 45.02 | 31.64 | 23.34 | 0.048 | 40 | 19 | 4 |
| 22 | V1 | 59.21 | 27.61 | 13.18 | 0.036 | 45 | 18 | 5 |
| 23 | V1 | 46.02 | 37.00 | 16.84 | 0.036 | 47 | 19 | 4 |
| 24 | V1 | 51.37 | 33.83 | 14.61 | 0.048 | 49 | 19* | 4* |

**Table S3:**
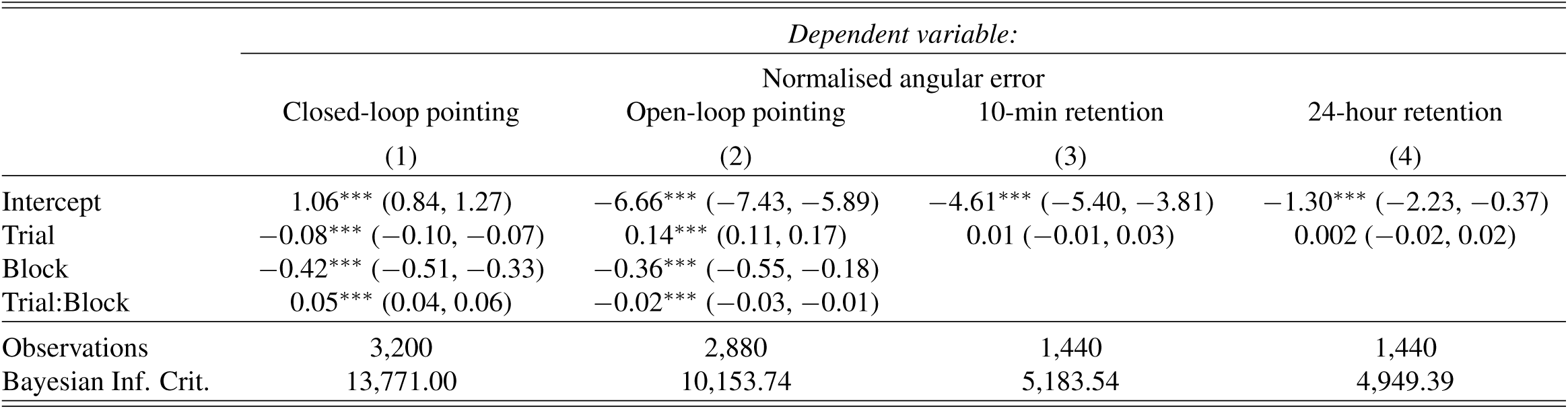
Experiment 1: Prism adaptation behaviour. All LMMs analysed the normalised pointing error as the dependent variable (i.e. trial endpoint errors *minus* mean baseline error). Model (1) assesses the reduction of CLP errors throughout prism exposure (blocks E1-6), while model (2) captures the development of an after-effect on OLP trials (blocks AE1-6). Models (3) and assess the persistence (intercept) and stability (main effect of Trial) of the after-effect (OLP) at the 10-minutes and 24-hours retention intervals. *^∗^*p*<*0.1; *^∗∗^*p*<*0.05; *^∗∗∗^*p*<*0.01 (all two-tailed).

**Table S4:** Experiment 1: Older participants show stronger long-term retention. The LMMs reported in this table examine the relationship between age and prism adaptation memory. The normalised pointing error was the dependent variable (i.e. trial endpoint errors *minus* mean baseline error). Models (1), (2) and (3) examine the relationship between age and prism after-effect at the end of adaptation (block AE6), and at the 10-minutes and 24-hour retention time points, respectively. Only 24-hour retention was related to age, such that older participants showed a larger (more negative) after-effect (AE). The next three models assess the robustness of this result when controlling for the average AE at the end of adaptation (model 4), the average AE at the 10- minutes retention interval (model 5), and the average movement duration on CLP trials during prism exposure (model 6). The relationship between age and long-term adaptation memory survived controlling for all three factors, confirming that it was not an artefact of older participants adapting to a greater extent on the first day or pointing more slowly. *^∗^*p*<*0.1; *^∗∗^*p*<*0.05; *^∗∗∗^*p*<*0.01 (all two-tailed).

|  | *Dependent variable:* |  |  |  |  |  |
| --- | --- | --- | --- | --- | --- | --- |
|  | End of PA<br>(1) | 10-min retention<br>(2) | 24-hour retention<br>(3) | Normalised angular error<br>24-hour retention [c:AE6]<br>(4) | 24-hour retention [controlling:10-min]<br>(5) | 24-hour retention [c:mvtDuration]<br>(6) |
| Intercept | −7.36*** (−8.46, −6.26) | −4.61*** (−5.39, −3.82) | −1.30*** (−2.16, −0.44) | −1.30*** (−2.16, −0.44) | −1.30*** (−2.16, −0.44) | −0.07 (−3.36, 3.21) |
| Trial | 0.10*** (0.07, 0.14) | 0.01 (−0.01, 0.03) | 0.002 (−0.02, 0.02) | 0.002 (−0.02, 0.02) | 0.002 (−0.02, 0.02) | 0.002 (−0.02, 0.02) |
| Age | 0.03 (−0.11, 0.17) | 0.05 (−0.05, 0.15) | −0.12** (−0.23, −0.02) | −0.13** (−0.24, −0.02) | −0.13** (−0.24, −0.02) | −0.12** (−0.23, −0.01) |
| Age:Trial | 0.0005 (−0.004, 0.01) | 0.002* (−0.0003, 0.004) | 0.001 (−0.002, 0.004) | 0.001 (−0.002, 0.004) | 0.001 (−0.002, 0.004) | 0.001 (−0.002, 0.004) |
| mean AE (end of PA) |  |  |  | 0.11 (−0.14, 0.37) |  |  |
| mean AE (10-min) |  |  |  |  | 0.03 (−0.33, 0.40) |  |
| Movement duration |  |  |  |  |  | −0.003 (−0.01, 0.004) |
| Observations | 480 | 1,440 | 1,440 | 1,440 | 1,440 | 1,440 |
| Bayesian Inf. Crit. | 1,730.39 | 5,194.85 | 4,957.99 | 4,964.52 | 4,965.23 | 4,964.71 |

**Table S5:** Experiment 1: Older participants have a higher excitation:inhibition ratio in sensorimotor cortex. The linear regressions reported in this table examine the relationship between age and metabolite concentration within the motor (labelled “M1”) and occipital (labelled “V1”) cortex voxels. All models controlled for the fraction of grey and white matter within the MRS voxel, and included the MRS measure as the dependent variable. Model (1) shows the predicted significant positive relationship between age and E:I ratio (Glx:GABA). Models (2) and (3) decompose this relationship into its GABA and Glx constituents respectively. They highlight that the age-related increase in E:I was mainly due to a loss of GABA-ergic inhibition. The final three models show a qualitatively similar, though not significant, pattern within the bilateral occipital cortex. *^∗^*p*<*0.1; *^∗∗^*p*<*0.05; *^∗∗∗^*p*<*0.01 (all two-tailed).

|  | *Dependent variable:* |  |  |  |  |  |
| --- | --- | --- | --- | --- | --- | --- |
|  | M1 E:I | M1 GABA | M1 Glx | V1 E:I | V1 GABA | V1 Glx |
|  | (1) | (2) | (3) | (4) | (5) | (6) |
| Intercept | −0.00 (−0.41, 0.41) | 0.00 (−0.14, 0.14) | 0.00 (−0.39, 0.39) | −0.22 (−0.73, 0.30) | 0.08 (−0.12, 0.27) | −0.34** (−0.63, −0.05) |
| Age | 0.08* (0.005, 0.15) | −0.03** (−0.06, −0.01) | −0.03 (−0.11, 0.05) | 0.05 (−0.02, 0.12) | −0.02 (−0.04, 0.004) | 0.005 (−0.04, 0.05) |
| Glx |  | 0.04 (−0.13, 0.21) |  |  | 0.28* (0.003, 0.56) |  |
| GABA |  |  | 0.31 (−0.98, 1.61) |  |  | 0.93* (0.01, 1.85) |
| GM | 0.15 (−0.04, 0.34) | −0.07* (−0.14, −0.0003) | −0.07 (−0.27, 0.13) | −0.10 (−0.24, 0.05) | 0.05* (−0.003, 0.10) | −0.12** (−0.20, −0.03) |
| WM | 0.04 (−0.05, 0.13) | −0.03 (−0.07, 0.01) | −0.12** (−0.22, −0.03) | −0.11 (−0.24, 0.02) | 0.06 (−0.01, 0.14) | −0.22*** (−0.29, −0.15) |
| Observations | 22 | 22 | 22 | 16 | 16 | 16 |
| Adjusted R<sup>2</sup> | 0.06 | 0.20 | 0.25 | 0.40 | 0.48 | 0.77 |

**Table S6:** Experiment 1: Higher sensorimotor cortex excitation:inhibition ratio is associated with greater 24-hour retention. The LMMs reported in this table examine the relationship between M1 and V1 neurochemistry and the magnitude of the AE at 24-hours. All models controlled for the fraction of grey and white matter within the MRS voxel. Model (1) shows that individuals with higher M1 E:I had a larger (more negative) AE at 24-hours. Model (2) decomposes this relationship into its GABA and Glx constituents respectively, highlighting that GABA but not Glx drives the previous relationship. Models (3) and (4) show that these findings were robust to controlling for the average movement duration on CLP trials during prism exposure. Finally, models to (8) reproduce the same set of analyses using MRS data from the anatomical control voxel (V1). No relationship between neurochemistry and long-term adaptation memory was observed in the V1 voxel. *^∗^*p*<*0.1; *^∗∗^*p*<*0.05; *^∗∗∗^*p*<*0.01 (all two-tailed).

|  | *Dependent variable:* |  |  |  |  |  |  |  |
| --- | --- | --- | --- | --- | --- | --- | --- | --- |
|  | Normalised angular error |  |  |  |  |  |  |  |
|  | M1 E:I<br>(1) | M1 GABA and Glx<br>(2) | M1 E:I [c:mvtDuration]<br>(3) | M1 GABA and Glx [c:mvtDuration]<br>(4) | V1 E:I<br>(5) | V1 GABA and Glx<br>(6) | V1 E:I [c:mvtDuration]<br>(7) | V1 GABA and Glx [c:mvtDuration]<br>(8) |
| Intercept | −0.79** (−1.53, −0.05) | −0.79** (−1.56, −0.03) | 2.73 (−1.41, 6.87) | 1.51 (−2.86, 5.87) | −0.44 (−2.07, 1.20) | 0.08 (−1.75, 1.91) | 10.95*** (4.23, 17.67) | 11.12*** (4.14, 18.11) |
| Trial | 0.01 (−0.03, 0.04) | 0.01 (−0.03, 0.04) | 0.01 (−0.03, 0.04) | 0.01 (−0.03, 0.04) | −0.001 (−0.04, 0.04) | −0.003 (−0.04, 0.04) | −0.001 (−0.04, 0.04) | −0.003 (−0.04, 0.04) |
| EI | −2.07*** (−2.82, −1.32) |  | −2.07*** (−2.77, −1.37) |  | 0.21 (−1.41, 1.82) |  | 1.02 (−0.35, 2.39) |  |
| GABA |  | 5.59*** (3.42, 7.76) |  | 5.58*** (3.49, 7.67) |  | −1.51 (−6.62, 3.60) |  | −3.32 (−7.63, 0.99) |
| Glx |  | 0.003 (−0.87, 0.88) |  | −0.15 (−1.04, 0.74) |  | 1.70 (−1.24, 4.63) |  | 0.48 (−1.97, 2.94) |
| GM | 0.24** (0.03, 0.45) | 0.29*** (0.09, 0.50) | 0.15 (−0.07, 0.37) | 0.23** (0.01, 0.45) | 0.28 (−0.11, 0.67) | 0.44* (−0.05, 0.94) | 0.20 (−0.10, 0.50) | 0.23 (−0.17, 0.64) |
| WM | −0.01 (−0.15, 0.12) | 0.11 (−0.05, 0.27) | −0.03 (−0.16, 0.10) | 0.08 (−0.08, 0.24) | 0.30 (−0.08, 0.69) | 0.65* (−0.06, 1.36) | 0.18 (−0.12, 0.48) | 0.18 (−0.45, 0.80) |
| Trial:EI | −0.01 (−0.04, 0.02) |  | −0.01 (−0.04, 0.02) |  | 0.01 (−0.02, 0.05) |  | 0.01 (−0.02, 0.05) |  |
| Trial:GABA |  | 0.01 (−0.08, 0.10) |  | 0.01 (−0.08, 0.10) |  | −0.03 (−0.13, 0.08) |  | −0.03 (−0.13, 0.08) |
| Trial:Glx |  | −0.004 (−0.03, 0.03) |  | −0.004 (−0.03, 0.03) |  | 0.01 (−0.03, 0.05) |  | 0.01 (−0.03, 0.05) |
| Movement duration |  |  | −0.01* (−0.02, 0.001) | −0.01 (−0.02, 0.01) |  |  | −0.03*** (−0.05, −0.01) | −0.03*** (−0.05, −0.01) |
| Observations | 990 | 990 | 990 | 990 | 720 | 720 | 720 | 720 |
| Bayesian Inf. Crit. | 3,359.62 | 3,371.64 | 3,363.89 | 3,377.51 | 2,479.94 | 2,491.97 | 2,477.90 | 2,490.77 |

**Table S7:** Experiment 1: Motor cortical GABA mediates the relationship between age and long-term adaptation memory. Model (1) shows a near-significant relationship between age and motor cortical E:I (*p* = 0.051, two-tailed; *p* = 0.025, one-tailed). Models (2) and (3) show that this relationship is driven by GABA (*p* = 0.013) and not Glx (*p* = 0.27). Model (4) shows that older age is associated with greater 24-hour retention (*p* = 0.02). Crucially, model (5) demonstrates that the association between age and 24-hour retention is no longer significant when accounting for M1 E:I. Further, model (6) shows that the mediation is specifically driven by GABA (*p* = 0.004) and not Glx (*p* = 0.78). Overall, these regression models provide support in favour of M1 GABA mediating the relationship between age and long-term retention, which was subsequently assessed formally. The mediation analysis indicates a significant effect of M1 GABA (*ab*_1_ = −0.50, 95%*CI* : [−1.36*, −*0.14]*, p* = 0.01) but not M1 Glx (*ab*_2_ = 0.02, 95%*CI* : [−0.09, 0.31]*, p* = 0.73; see Fig. 4).*^∗^*p*<*0.1; *^∗∗^*p*<*0.05; *^∗∗∗^*p*<*0.01 (all two-tailed).

|  | *Dependent variable:* |  |  |  |  |  |
| --- | --- | --- | --- | --- | --- | --- |
|  | M<sub>1</sub> (M1 E:I) | M<sub>2</sub> (M1 GABA) | M<sub>3</sub> (M1 Glx) |  | Y (AE24hrs) |  |
|  | (1) | (2) | (3) | (4) | (5) | (6) |
| X (age) | 0.66* (0.31) | −0.78** (0.28) | −0.31 (0.27) | −0.76** (0.29) | −0.35 (0.24) | −0.28 (0.28) |
| M<sub>1</sub> (M1 E:I) |  |  |  |  | −0.63*** (0.16) |  |
| M<sub>2</sub> (M1 GABA) |  |  |  |  |  | 0.65*** (0.20) |
| M<sub>3</sub> (M1 Glx) |  |  |  |  |  | −0.06 (0.20) |
| C<sub>1</sub> (GM) | 0.54 (0.35) | −0.68** (0.31) | −0.31 (0.30) | −0.40 (0.32) | −0.07 (0.25) | 0.02 (0.29) |
| C<sub>2</sub> (WM) | 0.20 (0.26) | −0.53** (0.23) | −0.70*** (0.22) | −0.32 (0.23) | −0.19 (0.18) | −0.01 (0.25) |
| Observations | 22 | 22 | 22 | 22 | 22 | 22 |
| Adjusted R<sup>2</sup> | 0.06 | 0.24 | 0.28 | 0.22 | 0.56 | 0.48 |
| F Statistic | 1.46 (df = 3; 18) | 3.16* (df = 3; 18) | 3.72** (df = 3; 18) | 2.99* (df = 3; 18) | 7.81*** (df = 4; 17) | 4.88*** (df = 5; 16) |

**Table S8:** Experiment 1: Mediation analysis controlling for average CLP duration. This table presents the results of the mediation analysis, controlling for the average reaching movement duration on CLP trials during prism exposure. Overall, this table indicates that the results presented in Table S7 are unchanged: M1 GABA, but not Glx, mediates the relationship between age and 24-hour retention.*^∗^*p*<*0.1; *^∗∗^*p*<*0.05; *^∗∗∗^*p*<*0.01 (all two-tailed).

|  | *Dependent variable:* |  |  |  |  |  |
| --- | --- | --- | --- | --- | --- | --- |
|  | M<sub>1</sub> (M1 E:I) | M<sub>2</sub> (M1 GABA) | M<sub>3</sub> (M1 Glx) |  | Y (AE24hrs) |  |
|  | (1) | (2) | (3) | (4) | (5) | (6) |
| X (age) | 0.66* (0.31) | −0.78** (0.28) | −0.31 (0.27) | −0.68** (0.29) | −0.34 (0.24) | −0.28 (0.29) |
| M<sub>1</sub> (M1 E:I) |  |  |  |  | −0.60*** (0.17) |  |
| M<sub>2</sub> (M1 GABA) |  |  |  |  |  | 0.66** (0.23) |
| M<sub>3</sub> (M1 Glx) |  |  |  |  |  | −0.05 (0.21) |
| C<sub>1</sub> (GM) | 0.54 (0.35) | −0.68** (0.31) | −0.31 (0.30) | −0.31 (0.32) | −0.04 (0.26) | 0.02 (0.30) |
| C<sub>2</sub> (WM) | 0.20 (0.26) | −0.53** (0.23) | −0.70*** (0.22) | −0.27 (0.23) | −0.18 (0.18) | −0.01 (0.27) |
| C<sub>3</sub> (Mvt duration) |  |  |  | −0.23 (0.20) | −0.09 (0.16) | 0.02 (0.20) |
| Observations | 22 | 22 | 22 | 22 | 22 | 22 |
| Adjusted R<sup>2</sup> | 0.06 | 0.24 | 0.28 | 0.24 | 0.55 | 0.45 |
| F Statistic | 1.46 (df = 3; 18) | 3.16* (df = 3; 18) | 3.72** (df = 3; 18) | 2.63* (df = 4; 17) | 6.08*** (df = 5; 16) | 3.81** (df = 6; 15) |

**Table S9:** Experiment 2: Association between a-tDCS, M1 neurochemistry, and adaptation memory. All linear mixed-effect models use the normalised pointing error (at 10-mins or 24-hours post-PA) as the dependent variable. Model (1) assesses the effect of left M1 a-tDCS on the after-effect at 10-min. Model (2) assesses the interaction of left M1 E:I (Glx:GABA) with this effect. Model (3) decomposes the individual interaction of GABA and Glx with the behavioural effect of a-tDCS effect on short-term retention. Models (4), (5), (6) assess the same effects at the long-term retention time point (24-hours). The most important finding here is that M1 E:I significantly interacted with the effect of a-tDCS on long-term retention (a-tDCS:EI in Model 5), which could be decomposed into opposite interactions with GABA and Glx (a-tDCS:GABA and a-tDCS:Glx in Model 6). All models including MRS data also controlled for the fraction of grey and white matter within the voxel. *^∗^*p*<*0.1; *^∗∗^*p*<*0.05; *^∗∗∗^*p*<*0.01 (all two-tailed).

|  | *Dependent variable:* |  |  |  |  |  |
| --- | --- | --- | --- | --- | --- | --- |
|  | Normalised angular error |  |  |  |  |  |
|  | a-tDCS:10min<br>(1) | a-tDCS:EI (10min)<br>(2) | a-tDCS:GABA/Glx (10min)<br>(3) | a-tDCS:24-hour<br>(4) | a-tDCS:EI (24hrs)<br>(5) | a-tDCS:GABA/Glx (24hrs)<br>(6) |
| Intercept | −4.7*** (−5.5, −3.8) | −5.0*** (−6.1, −4.0) | −5.0*** (−6.1, −4.0) | −1.0** (−1.8, −0.2) | −1.1*** (−1.7, −0.4) | −1.1*** (−1.7, −0.5) |
| Trial | −0.003 (−0.02, 0.01) | 0.003 (−0.02, 0.02) | 0.003 (−0.02, 0.02) | −0.001 (−0.02, 0.02) | −0.001 (−0.03, 0.03) | −0.001 (−0.03, 0.03) |
| GM |  | 0.1 (−0.1, 0.3) | 0.1 (−0.1, 0.3) |  | 0.2*** (0.1, 0.4) | 0.2*** (0.1, 0.4) |
| WM |  | −0.1 (−0.2, 0.04) | −0.04 (−0.2, 0.1) |  | 0.1 (−0.05, 0.2) | 0.1 (−0.1, 0.2) |
| a-tDCS | 0.1 (−0.5, 0.6) | 0.1 (−0.5, 0.7) | 0.1 (−0.5, 0.7) | −0.5 (−1.3, 0.2) | 0.04 (−0.6, 0.7) | 0.04 (−0.6, 0.7) |
| E:I |  | −0.4 (−1.5, 0.6) |  |  | −0.9*** (−1.5, −0.3) |  |
| GABA |  |  | 0.6 (−2.4, 3.6) |  |  | 2.8*** (1.0, 4.6) |
| Glx |  |  | −0.1 (−1.2, 1.0) |  |  | −0.3 (−1.0, 0.5) |
| Trial:a-tDCS | −0.01 (−0.02, 0.01) | 0.000 (−0.01, 0.01) | 0.000 (−0.01, 0.01) | −0.01 (−0.02, 0.005) | −0.01 (−0.02, 0.01) | −0.01 (−0.02, 0.01) |
| Trial:EI |  | 0.01 (−0.01, 0.03) |  |  | 0.02 (−0.01, 0.05) |  |
| Trial:GABA |  |  | −0.03 (−0.1, 0.03) |  |  | −0.04 (−0.1, 0.04) |
| Trial:Glx |  |  | 0.01 (−0.01, 0.03) |  |  | 0.004 (−0.02, 0.03) |
| a-tDCS:EI |  | 0.6* (−0.03, 1.2) |  |  | 0.8** (0.1, 1.4) |  |
| a-tDCS:GABA |  |  | −1.2 (−3.0, 0.6) |  |  | −1.5* (−3.3, 0.2) |
| a-tDCS:Glx |  |  | 0.3 (−0.3, 0.8) |  |  | 0.7** (0.2, 1.3) |
| Trial:a-tDCS:EI |  | 0.01 (−0.01, 0.02) |  |  | 0.01** (0.002, 0.02) |  |
| Trial:a-tDCS:GABA |  |  | −0.01 (−0.1, 0.03) |  |  | −0.03 (−0.1, 0.01) |
| Trial:a-tDCS:Glx |  |  | 0.01 (−0.003, 0.02) |  |  | 0.004 (−0.01, 0.01) |
| Observations | 2,250 | 1,440 | 1,440 | 2,250 | 1,440 | 1,440 |
| Bayesian Inf. Crit. | 7,470.8 | 4,830.3 | 4,858.5 | 8,378.3 | 5,400.6 | 5,425.6 |

Note:
\*p<0.1; \*\*p<0.05; \*\*\*p<0.01

**Table S10:** Experiment 2: Association between a-tDCS, M1 neurochemistry, and adaptation memory while controlling for average movement speed during prism exposure. All linear mixed-effect models use the normalised pointing error (at 10-min or 24-hours post-PA) as the dependent variable, and control for the average movement duration of the CLP trials of the prism exposure. Model (1) assesses the moderating influence of left M1 E:I (Glx:GABA) on the behavioural effect of a-tDCS on the AE at 10-minutes. Model (2) decomposes the individual interaction of GABA and Glx with the behavioural effect of a-tDCS effect on short-term retention. Models (4) and (5) assess the same effects at the long-term retention time point (24-hours). There was a significant interaction between a-tDCS and E:I at the 24-hours retention time point. All models also controlled for the fraction of grey and white matter within the voxel. *^∗^*p*<*0.1; *^∗∗^*p*<*0.05; *^∗∗∗^*p*<*0.01 (all two-tailed).

|  | *Dependent variable:* |  |  |  |
| --- | --- | --- | --- | --- |
|  | Normalised angular error |  |  |  |
|  | a-tDCS:EI (10min) | a-tDCS:GABA/Glx (10min) | a-tDCS:EI (24hrs) | a-tDCS:GABA/Glx (24hrs) |
|  | (1) | (2) | (3) | (4) |
| Intercept | −1.9 (−7.6, 3.8) | −1.9 (−9.4, 5.6) | −4.3* (−9.0, 0.4) | −7.4** (−13.1, −1.7) |
| Trial | 0.003 (−0.02, 0.02) | 0.003 (−0.02, 0.02) | −0.001 (−0.03, 0.03) | −0.001 (−0.03, 0.03) |
| Mvt Duration | −0.01 (−0.02, 0.005) | −0.01 (−0.02, 0.01) | 0.01 (−0.003, 0.01) | 0.01** (0.001, 0.02) |
| GM | 0.01 (−0.2, 0.3) | 0.01 (−0.3, 0.3) | 0.3*** (0.1, 0.5) | 0.4*** (0.2, 0.6) |
| WM | −0.1* (−0.2, 0.02) | −0.1 (−0.3, 0.1) | 0.1 (−0.03, 0.2) | 0.1* (−0.01, 0.3) |
| a-tDCS | 0.1 (−0.5, 0.7) | 0.1 (−0.5, 0.7) | 0.04 (−0.6, 0.7) | 0.04 (−0.6, 0.7) |
| EI | −0.3 (−1.3, 0.7) |  | −1.0*** (−1.7, −0.4) |  |
| GABA |  | 0.2 (−2.9, 3.2) |  | 3.6*** (1.7, 5.6) |
| Glx |  | −0.4 (−1.6, 0.8) |  | 0.2 (−0.7, 1.0) |
| Trial:a-tDCS | 0.000 (−0.01, 0.01) | 0.000 (−0.01, 0.01) | −0.01 (−0.02, 0.01) | −0.01 (−0.02, 0.01) |
| Trial:EI | 0.01 (−0.01, 0.03) |  | 0.02 (−0.01, 0.05) |  |
| Trial:GABA |  | −0.03 (−0.1, 0.03) |  | −0.04 (−0.1, 0.04) |
| Trial:Glx |  | 0.01 (−0.01, 0.03) |  | 0.004 (−0.02, 0.03) |
| a-tDCS:EI | 0.6* (−0.03, 1.2) |  | 0.8** (0.1, 1.4) |  |
| a-tDCS:GABA |  | −1.2 (−3.0, 0.6) |  | −1.5* (−3.3, 0.2) |
| a-tDCS:Glx |  | 0.3 (−0.3, 0.8) |  | 0.7** (0.2, 1.3) |
| Trial:a-tDCS:EI | 0.01 (−0.01, 0.02) |  | 0.01** (0.002, 0.02) |  |
| Trial:a-tDCS:GABA |  | −0.01 (−0.1, 0.03) |  | −0.03 (−0.1, 0.01) |
| Trial:a-tDCS:Glx |  | 0.01 (−0.003, 0.02) |  | 0.004 (−0.01, 0.01) |
| Observations | 1,440 | 1,440 | 1,440 | 1,440 |
| Bayesian Inf. Crit. | 4,836.5 | 4,865.2 | 5,406.9 | 5,430.6 |

Note:
\*p<0.1; \*\*p<0.05; \*\*\*p<0.01

**Table S11:** Experiment 2: Association between a-tDCS, V1 neurochemistry, and adaptation memory. All linear mixed-effect models use the normalised pointing error (at 10-min or 24-hours post-PA) as the dependent variable. Model (1) assesses the moderating influence of left M1 E:I (Glx:GABA) on the behavioural effect of a-tDCS on the AE at 10-minutes. Model (2) decomposes the individual interaction of GABA and Glx with the behavioural effect of a-tDCS effect on short-term retention. Models (4) and (5) assess the same effects at the long-term retention time point (24-hours). There was no significant interaction between a-tDCS and E:I at either of the two retention time points. All models also controlled for the fraction of grey and white matter within the voxel. *^∗^*p*<*0.1; *^∗∗^*p*<*0.05; *^∗∗∗^*p*<*0.01 (all two-tailed).

|  | *Dependent variable:* |  |  |  |
| --- | --- | --- | --- | --- |
|  | Normalised angular error |  |  |  |
|  | a-tDCS:EI (10min) | a-tDCS:GABA/Glx (10min) | a-tDCS:EI (24hrs) | a-tDCS:GABA/Glx (24hrs) |
|  | (1) | (2) | (3) | (4) |
| Intercept | −4.1*** (−5.4, −2.8) | −3.3*** (−4.7, −1.9) | −2.0*** (−3.2, −0.9) | −1.3** (−2.4, −0.2) |
| Trial | 0.004 (−0.02, 0.03) | 0.004 (−0.02, 0.03) | −0.01 (−0.04, 0.03) | −0.01 (−0.05, 0.02) |
| GM | 0.2 (−0.1, 0.4) | 0.3** (0.04, 0.5) | −0.2* (−0.5, 0.04) | −0.3*** (−0.6, −0.1) |
| WM | 0.2 (−0.1, 0.4) | 0.5*** (0.1, 0.8) | −0.3** (−0.5, −0.01) | −0.1 (−0.4, 0.2) |
| a-tDCS | 0.3 (−0.5, 1.0) | 0.2 (−0.5, 0.8) | 0.4 (−0.5, 1.2) | 0.3 (−0.6, 1.1) |
| EI | 0.3 (−0.9, 1.5) |  | −1.3** (−2.4, −0.2) |  |
| GABA |  | −2.6 (−6.5, 1.3) |  | 4.9*** (1.9, 7.8) |
| Glx |  | 0.8 (−1.0, 2.6) |  | 0.4 (−1.2, 1.9) |
| Trial:a-tDCS | −0.001 (−0.02, 0.02) | −0.002 (−0.02, 0.02) | −0.002 (−0.02, 0.01) | −0.003 (−0.02, 0.01) |
| Trial:EI | −0.01 (−0.03, 0.02) |  | 0.005 (−0.03, 0.04) |  |
| Trial:GABA |  | 0.01 (−0.1, 0.1) |  | −0.004 (−0.1, 0.1) |
| Trial:Glx |  | −0.000 (−0.03, 0.03) |  | 0.01 (−0.03, 0.05) |
| a-tDCS:EI | −0.4 (−1.1, 0.3) |  | 0.4 (−0.4, 1.1) |  |
| a-tDCS:GABA |  | 2.6** (0.5, 4.7) |  | −0.6 (−3.2, 1.9) |
| a-tDCS:Glx |  | 0.5 (−0.2, 1.3) |  | 0.5 (−0.4, 1.5) |
| Trial:a-tDCS:EI | −0.005 (−0.02, 0.01) |  | 0.005 (−0.01, 0.02) |  |
| Trial:a-tDCS:GABA |  | 0.03 (−0.03, 0.1) |  | −0.000 (−0.05, 0.05) |
| Trial:a-tDCS:Glx |  | 0.01 (−0.01, 0.03) |  | 0.01 (−0.01, 0.03) |
| Observations | 1,080 | 1,080 | 1,080 | 1,080 |
| Bayesian Inf. Crit. | 3,639.9 | 3,661.8 | 4,148.1 | 4,170.5 |

Note:
\*p<0.1; \*\*p<0.05; \*\*\*p<0.01

## Notes

### Competing Interest Statement

The authors have declared no competing interest.

## References

1. Krampe, R. T. Aging, expertise and fine motor movement. Neuroscience Biobehavioral Reviews 26, 769 – 776 (2002). URL http://www.sciencedirect.com/science/article/pii/S0149763402000647.

2. Hunter, S. K., Pereira, H. M. & Keenan, K. G. The aging neuromuscular system and motor performance. Journal of Applied Physiology 121, 982–995 (2016). URL https://doi.org/10.1152/japplphysiol.00475.2016.

3. Bedard, A.-C. et al. The development of selective inhibitory control across the life span. Developmental Neuropsychology 21, 93–111 (2002). URL https://doi.org/10.1207/S15326942DN2101_5.

4. Jimnez-Jimnez, F. J. et al. Influence of age and gender in motor performance in healthy subjects. Journal of the Neurological Sciences 302, 72 – 80 (2011). URL http://www.sciencedirect.com/science/article/pii/S0022510X10005691.

5. Frontera, W. R., et al. Aging of skeletal muscle: a 12-yr longitudinal study. Journal of Applied Physiology 88, 1321–1326 (2000). URL https://doi.org/10.1152/jappl.2000.88.4.1321.

6. Serrien, D. J., Swinnen, S. P. & Stelmach, G. E. Age-Related Deterioration of Coordinated Interlimb Behavior. The Journals of Gerontology: Series B 55, P295–P303 (2000). URL https://doi.org/10.1093/geronb/55.5.P295.

7. Leeson, G. W. The growth, ageing and urbanisation of our world. Journal of Population Ageing 11, 107–115 (2018). URL https://doi.org/10.1007/s12062-018-9225-7.

8. Dayan, E. & Cohen, L. Neuroplasticity subserving motor skill learning. Neuron 72, 443 – 454 (2011). URL http://www.sciencedirect.com/science/article/pii/S0896627311009184.

9. Sampaio-Baptista, C., Sanders, Z.-B. & Johansen-Berg, H. Structural plasticity in adulthood with motor learning and stroke rehabilitation. Annual Review of Neuroscience 41, 25–40 (2018). URL https://doi.org/10.1146/annurev-neuro-080317-062015.

10. Rozycka, A. & Liguz-Lecznar, M. The space where aging acts: focus on the GABAergic synapse. Aging Cell 16, 634–643 (2017). URL https://onlinelibrary.wiley.com/doi/abs/10.1111/acel.12605.

11. McNeil, C. J. & Rice, C. L. Neuromuscular adaptations to healthy aging. Applied Physiology, Nutrition, and Metabolism 43, 1158–1165 (2018). URL https://doi.org/10.1139/apnm-2018-0327.

12. Burke, S. N. & Barnes, C. A. Neural plasticity in the ageing brain. Nature Reviews Neuroscience 7, 30–40 (2006). URL https://doi.org/10.1038/nrn1809.

13. Rogasch, N. C., Dartnall, T. J., Cirillo, J., Nordstrom, M. A. & Semmler, J. G. Corticomo-tor plasticity and learning of a ballistic thumb training task are diminished in older adults. Journal of Applied Physiology 107, 1874–1883 (2009). URL https://doi.org/10.1152/japplphysiol.00443.2009.

14. Freitas, C., Farzan, F. & Pascual-Leone, A. Assessing brain plasticity across the lifespan with transcranial magnetic stimulation: why, how, and what is the ultimate goal? Frontiers in Neuroscience 7, 42 (2013). URL https://www.frontiersin.org/article/10.3389/fnins.2013.00042.

15. Bhandari, A. et al. A meta-analysis of the effects of aging on motor cortex neurophysiology assessed by transcranial magnetic stimulation. Clinical Neurophysiology 127, 2834–2845 (2016). URL http://www.sciencedirect.com/science/article/pii/S1388245716304266.

16. David-Jrgens, M. & Dinse, H. R. Effects of Aging on Paired-Pulse Behavior of Rat Somatosensory Cortical Neurons. Cerebral Cortex 20, 1208–1216 (2009). URL https://doi.org/10.1093/cercor/bhp185.

17. Schmidt, S., Redecker, C., Bruehl, C. & Witte, O. Age-related decline of functional inhibition in rat cortex. Neurobiology of Aging 31, 504 – 511 (2010). URL http://www.sciencedirect.com/science/article/pii/S0197458008001309.

18. Peinemann, A., Lehner, C., Conrad, B. & Siebner, H. R. Age-related decrease in paired-pulse intracortical inhibition in the human primary motor cortex. Neuroscience Letters 313, 33– 36 (2001). URL https://www.sciencedirect.com/science/article/pii/S030439400102239X.

19. Oliviero, A. et al. Effects of aging on motor cortex excitability. Neuroscience Research 55, 74–77 (2006). URL https://www.sciencedirect.com/science/article/pii/S0168010206000289.

20. Lenz, M. et al. Increased excitability of somatosensory cortex in aged humans is associated with impaired tactile acuity. Journal of Neuroscience 32, 1811–1816 (2012). URL https://www.jneurosci.org/content/32/5/1811.

21. Cheng, C.-H. & Lin, Y.-Y. Aging-related decline in somatosensory inhibition of the human cerebral cortex. Experimental Brain Research 226, 145–152 (2013). URL https://doi.org/10.1007/s00221-013-3420-9.

22. Gao, F. et al. Edited magnetic resonance spectroscopy detects an age-related decline in brain gaba levels. NeuroImage 78, 75–82 (2013). URL https://www.sciencedirect.com/science/article/pii/S105381191300339X.

23. Heise, K.-F. et al. The aging motor system as a model for plastic changes of gaba-mediated intracortical inhibition and their behavioral relevance. Journal of Neuroscience 33, 9039– 9049 (2013). URL https://www.jneurosci.org/content/33/21/9039.

24. Levin, O., Fujiyama, H., Boisgontier, M. P., Swinnen, S. P. & Summers, J. J. Aging and motor inhibition: A converging perspective provided by brain stimulation and imaging approaches. Neuroscience Biobehavioral Reviews 43, 100 – 117 (2014). URL http://www.sciencedirect.com/science/article/pii/S0149763414000852.

25. Mooney, R. A., Cirillo, J. & Byblow, W. D. GABA and primary motor cortex inhibition in young and older adults: a multimodal reliability study. Journal of Neurophysiology 118, 425–433 (2017). URL https://doi.org/10.1152/jn.00199.2017.

26. Hermans, L. et al. Brain GABA levels are associated with inhibitory control deficits in older adults. Journal of Neuroscience 38, 7844–7851 (2018). URL https://www.jneurosci.org/content/38/36/7844.

27. Marenco, S. et al. Role of gamma-amino-butyric acid in the dorsal anterior cingulate in age-associated changes in cognition. Neuropsychopharmacology 43, 2285–2291 (2018). URL https://doi.org/10.1038/s41386-018-0134-5.

28. Simmonite, M. et al. Age-related declines in occipital GABA are associated with reduced fluid processing ability. Academic Radiology 26, 1053 – 1061 (2019). URL http://www.sciencedirect.com/science/article/pii/S1076633218304008.

29. Chamberlain, J. D. et al. GABA levels in ventral visual cortex decline with age and are associated with neural distinctiveness. Neurobiology of Aging (2021). URL https://www.sciencedirect.com/science/article/pii/S0197458021000683.

30. Kolasinski, J. et al. A mechanistic link from GABA to cortical architecture and perception. Current Biology 27, 1685–1691.e3 (2017). URL https://www.sciencedirect.com/science/article/pii/S0960982217305006.

31. Papegaaij, S., Taube, W., Hogenhout, M., Baudry, S. & Hortobgyi, T. Age-related decrease in motor cortical inhibition during standing under different sensory conditions. Frontiers in Aging Neuroscience 6, 126 (2014). URL https://www.frontiersin.org/article/10.3389/fnagi.2014.00126.

32. Swanson, C. W. & Fling, B. W. Associations between gait coordination, variability and motor cortex inhibition in young and older adults. Experimental Gerontology 113, 163 – 172 (2018). URL http://www.sciencedirect.com/science/article/pii/S0531556518304182.

33. Franklin, D. & Wolpert, D. Computational mechanisms of sensorimotor control. Neuron 72, 425–442 (2011). URL https://www.sciencedirect.com/science/article/pii/S0896627311008919.

34. Wolpert, D. M., Diedrichsen, J. & Flanagan, J. R. Principles of sensorimotor learning. Nature Reviews Neuroscience 12, 739–751 (2011). URL https://doi.org/10.1038/nrn3112.

35. Fernández-Ruiz, J., Hall, C., Vergara, P. & Díaz, R. Prism adaptation in normal aging: slower adaptation rate and larger aftereffect. Cognitive Brain Research 9, 223–226 (2000). URL https://www.sciencedirect.com/science/article/pii/S0926641099000579.

36. Roller, C. A., Cohen, H. S., Kimball, K. T. & Bloomberg, J. J. Effects of normal aging on visuo-motor plasticity. Neurobiology of Aging 23, 117–123 (2002). URL https://www.sciencedirect.com/science/article/pii/S0197458001002640.

37. Buch, E. R., Young, S. & Contreras-Vidal, J. L. Visuomotor adaptation in normal aging. Learning & memory 10, 55–63 (2003). URL https://doi.org/10.1101/lm.50303.

38. Bock, O. Components of sensorimotor adaptation in young and elderly subjects. Experimental Brain Research 160, 259–263 (2005).

39. Hegele, M. & Heuer, H. Adaptation to a direction-dependent visuomotor gain in the young and elderly. Psychological Research PRPF 74, 21 (2008). URL https://doi.org/10.1007/s00426-008-0221-z.

40. Anguera, J. A., Reuter-Lorenz, P. A., Willingham, D. T. & Seidler, R. D. Failure to engage spatial working memory contributes to age-related declines in visuomotor learning. Journal of Cognitive Neuroscience 23, 11–25 (2011). URL https://doi.org/10.1162/jocn.2010.21451.

41. Huang, H. J. & Ahmed, A. A. Older adults learn less, but still reduce metabolic cost, during motor adaptation. Journal of Neurophysiology 111, 135–144 (2014). URL https://doi.org/10.1152/jn.00401.2013.

42. Nemanich, S. T. & Earhart, G. M. How do age and nature of the motor task influence visuomotor adaptation? Gait Posture 42, 564–568 (2015). URL https://www.sciencedirect.com/science/article/pii/S0966636215008498.

43. Panouille’res, M. T. N., Joundi, R. A., Brittain, J.-S. & Jenkinson, N. Reversing motor adaptation deficits in the ageing brain using non-invasive stimulation. The Journal of Physiology 593, 3645–3655 (2015). URL https://physoc.onlinelibrary.wiley.com/doi/abs/10.1113/JP270484.

44. Malone, L. A. & Bastian, A. J. Age-related forgetting in locomotor adaptation. Neurobiology of Learning and Memory 128, 1–6 (2016). URL https://www.sciencedirect.com/science/article/pii/S1074742715002051.

45. Vandevoorde, K. & Orban de Xivry, J.-J. Internal model recalibration does not deteriorate with age while motor adaptation does. Neurobiology of Aging 80, 138–153 (2019). URL https://www.sciencedirect.com/science/article/pii/S0197458019301058.

46. Wolpe, N. et al. Age-related reduction in motor adaptation: brain structural correlates and the role of explicit memory. Neurobiology of Aging 90, 13–23 (2020). URL https://www.sciencedirect.com/science/article/pii/S019745802030049X.

47. Galea, J. M., Vazquez, A., Pasricha, N., Orban de Xivry, J.-J. & Celnik, P. Dissociating the Roles of the Cerebellum and Motor Cortex during Adaptive Learning: The Motor Cortex Retains What the Cerebellum Learns. Cerebral Cortex 21, 1761–1770 (2010). URL https://doi.org/10.1093/cercor/bhq246.

48. O’Shea, J. et al. Induced sensorimotor cortex plasticity remediates chronic treatment-resistant visual neglect. eLife 6, e26602 (2017). URL https://doi.org/10.7554/eLife.26602.

49. von Helmholtz, H. Treatise on physiological optics, vol. 3 (Courier Corporation, 1867).

50. Stagg, C. J. et al. Polarity-sensitive modulation of cortical neurotransmitters by transcranial stimulation. Journal of Neuroscience 29, 5202–5206 (2009). URL https://www.jneurosci.org/content/29/16/5202. https://www.jneurosci.org/content/29/16/5202.full.pdf.

51. Kim, S., Stephenson, M. C., Morris, P. G. & Jackson, S. R. tDCS-induced alterations in GABA concentration within primary motor cortex predict motor learning and motor memory: A 7T magnetic resonance spectroscopy study. NeuroImage 99, 237–243 (2014). URL https://www.sciencedirect.com/science/article/pii/S1053811914004558.

52. Antonenko, D. et al. tDCS-induced modulation of GABA levels and resting-state functional connectivity in older adults. Journal of Neuroscience 37, 4065–4073 (2017). URL https://www.jneurosci.org/content/37/15/4065. https://www.jneurosci.org/content/37/15/4065.full.pdf.

53. Smith, M. et al. Menstrual cycle effects on cortical excitability. Neurology 53, 2069–2069 (1999). URL https://n.neurology.org/content/53/9/2069.

54. Epperson, C. N. et al. Cortical γ-aminobutyric acid levels across the menstrual cycle in healthy women and those with premenstrual dysphoric disorder: A proton magnetic resonance spectroscopy study. Archives of general psychiatry 59, 851–858 (2002). URL https://doi.org/10.1001/archpsyc.59.9.851.

55. Gordon, J. L. et al. Ovarian hormone fluctuation, neurosteroids, and HPA axis dysregulation in perimenopausal depression: A novel heuristic model. American Journal of Psychiatry 172, 227–236 (2015). URL https://doi.org/10.1176/appi.ajp.2014.14070918.

56. Inoue, M. et al. Three timescales in prism adaptation. Journal of Neurophysiology 113, 328–338 (2015). URL https://doi.org/10.1152/jn.00803.2013.

57. Marinescu, I. E., Lawlor, P. N. & Kording, K. P. Quasi-experimental causality in neuroscience and behavioural research. Nature Human Behaviour 2, 891–898 (2018). URL https://doi.org/10.1038/s41562-018-0466-5.

58. Barron, H. et al. Unmasking latent inhibitory connections in human cortex to reveal dormant cortical memories. Neuron 90, 191–203 (2016). URL https://www.sciencedirect.com/science/article/pii/S0896627316001689.

59. Fritsch, B. et al. Direct current stimulation promotes bdnf-dependent synaptic plasticity: Potential implications for motor learning. Neuron 66, 198–204 (2010). URL https://www.sciencedirect.com/science/article/pii/S0896627310002382.

60. Roig, M., Ritterband-Rosenbaum, A., Lundbye-Jensen, J. & Nielsen, J. B. Aging increases the susceptibility to motor memory interference and reduces off-line gains in motor skill learning. Neurobiology of Aging 35, 1892–1900 (2014). URL https://www.sciencedirect.com/science/article/pii/S0197458014002280.

61. Zimerman, M. et al. Neuroenhancement of the aging brain: Restoring skill acquisition in old subjects. Annals of Neurology 73, 10–15 (2013). URL https://onlinelibrary.wiley.com/doi/abs/10.1002/ana.23761.

62. Siebner, H. R. et al. Preconditioning of low-frequency repetitive transcranial magnetic stimulation with transcranial direct current stimulation: Evidence for homeostatic plasticity in the human motor cortex. Journal of Neuroscience 24, 3379–3385 (2004). URL https://www.jneurosci.org/content/24/13/3379. https://www.jneurosci.org/content/24/13/3379.full.pdf.

63. Lang, N. et al. Preconditioning with transcranial direct current stimulation sensitizes the motor cortex to rapid-rate transcranial magnetic stimulation and controls the direction of after-effects. Biological psychiatry 56, 634–639 (2004).

64. Nitsche, M. A. et al. Timing-dependent modulation of associative plasticity by general network excitability in the human motor cortex. Journal of Neuroscience 27, 3807–3812 (2007). URL https://www.jneurosci.org/content/27/14/3807. https://www.jneurosci.org/content/27/14/3807.full.pdf.

65. Krause, B., Mrquez-Ruiz, J. & Kadosh, R. C. The effect of transcranial direct current stimulation: a role for cortical excitation/inhibition balance? Frontiers in human neuroscience 7 (2013).

66. Karabanov, A. et al. Consensus paper: Probing homeostatic plasticity of human cortex with non-invasive transcranial brain stimulation. Brain Stimulation 8, 993–1006 (2015). URL http://www.sciencedirect.com/science/article/pii/S1935861X15010190.

67. Perez, C. C. Invisible women: Exposing data bias in a world designed for men (Random House, 2019).

68. Horvath, J., Carter, O. & Forte, J. Transcranial direct current stimulation: five important issues we aren’t discussing (but probably should be). Frontiers in Systems Neuroscience 8, 2 (2014). URL https://www.frontiersin.org/article/10.3389/fnsys.2014.00002.

69. Li, C.-S. R., Padoa-Schioppa, C. & Bizzi, E. Neuronal correlates of motor performance and motor learning in the primary motor cortex of monkeys adapting to an external force field. Neuron 30, 593–607 (2001). URL https://www.sciencedirect.com/science/article/pii/S0896627301003014.

70. Richardson, A. G. et al. Disruption of primary motor cortex before learning impairs memory of movement dynamics. Journal of Neuroscience 26, 12466–12470 (2006). URL https://www.jneurosci.org/content/26/48/12466. https://www.jneurosci.org/content/26/48/12466.full.pdf.

71. Hadipour-Niktarash, A., Lee, C. K., Desmond, J. E. & Shadmehr, R. Impairment of retention but not acquisition of a visuomotor skill throughtime-dependent disruption of primary motor cortex. Journal of Neuroscience 27, 13413–13419 (2007). URL https://www.jneurosci.org/content/27/49/13413.

72. Hunter, T., Sacco, P., Nitsche, M. A. & Turner, D. L. Modulation of internal model formation during force field-induced motor learning by anodal transcranial direct current stimulation of primary motor cortex. The Journal of Physiology 587, 2949–2961 (2009). URL https://physoc.onlinelibrary.wiley.com/doi/abs/10.1113/jphysiol.2009.169284.

73. Landi, S. M., Baguear, F. & Della-Maggiore, V. One week of motor adaptation induces structural changes in primary motor cortex that predict long-term memory one year later. Journal of Neuroscience 31, 11808–11813 (2011). URL https://www.jneurosci.org/content/31/33/11808

74. . .Stagg, C. J. et al. Local GABA concentration is related to network-level resting functional connectivity. eLife 3, e01465 (2014). URL https://doi.org/10.7554/eLife.01465.

75. Bachtiar, V., Near, J., Johansen-Berg, H. & Stagg, C. J. Modulation of GABA and resting state functional connectivity by transcranial direct current stimulation. eLife 4, e08789 (2015). URL https://doi.org/10.7554/eLife.08789.

76. Muthukumaraswamy, S. D., Edden, R. A., Jones, D. K., Swettenham, J. B. & Singh, K. D. Resting gaba concentration predicts peak gamma frequency and fmri amplitude in response to visual stimulation in humans. Proceedings of the National Academy of Sciences 106, 8356–8361 (2009). URL https://www.pnas.org/content/106/20/8356.

77. Buzsa’ki, G. & Schomburg, E. W. What does gamma coherence tell us about inter-regional neural communication? Nature Neuroscience 18, 484–489 (2015). URL https://doi.org/10.1038/nn.3952.

78. Kitago, T. & Krakauer, J. W. Chapter 8 - motor learning principles for neurorehabilitation. In Barnes, M. P. & Good, D. C. (eds.) Neurological Rehabilitation, vol. 110 of *Handbook* of Clinical Neurology, 93–103 (Elsevier, 2013). URL https://www.sciencedirect.com/science/article/pii/B9780444529015000083.

79. Galea, J. M., Mallia, E., Rothwell, J. & Diedrichsen, J. The dissociable effects of punishment and reward on motor learning. Nature Neuroscience 18, 597–602 (2015). URL https://doi.org/10.1038/nn.3956.

80. Quattrocchi, G., Greenwood, R., Rothwell, J. C., Galea, J. M. & Bestmann, S. Reward and punishment enhance motor adaptation in stroke. *Journal of Neurology*, Neurosurgery & Psychiatry 88, 730–736 (2017). URL https://jnnp.bmj.com/content/88/9/730.

81. Buch, E. R. et al. Effects of tdcs on motor learning and memory formation: A consensus and critical position paper. Clinical Neurophysiology 128, 589–603 (2017). URL https://www.sciencedirect.com/science/article/pii/S1388245717300263.

82. Chalavi, S. et al. The neurochemical basis of the contextual interference effect. Neurobiology of Aging 66, 85–96 (2018). URL https://www.sciencedirect.com/science/article/pii/S0197458018300575.

83. Pauwels, L. et al. Challenge to promote change: The neural basis of the contextual interference effect in young and older adults. Journal of Neuroscience 38, 3333–3345 (2018). URL https://www.jneurosci.org/content/38/13/3333.

84. Kording, K. P., Tenenbaum, J. B. & Shadmehr, R. The dynamics of memory as a consequence of optimal adaptation to a changing body. Nature Neuroscience 10, 779–786 (2007). URL https://doi.org/10.1038/nn1901.

85. Kapur, N. Paradoxical functional facilitation in brain-behaviour research: A critical review. Brain 119, 1775–1790 (1996). URL https://doi.org/10.1093/brain/119.5.1775.

86. Pitchaimuthu, K. et al. Occipital gaba levels in older adults and their relationship to visual perceptual suppression. Scientific Reports 7, 14231 (2017). URL https://doi.org/10.1038/s41598-017-14577-5.

87. Faul, F., Erdfelder, E., Lang, A.-G. & Buchner, A. G*power 3: A flexible statistical power analysis program for the social, behavioral, and biomedical sciences. Behavior Research Methods 39, 175–191 (2007). URL https://doi.org/10.3758/BF03193146.

88. Kleiner, M., Brainard, D. & Pelli, D. What’s new in psychtoolbox-3? (2007).

89. Redding, G. M. & Wallace, B. Adaptive spatial alignment and strategic perceptual-motor control. Journal of Experimental Psychology: Human Perception and Performance 22, 379 (1996). URL https://doi.org/10.1037/0096-1523.22.2.379.

90. Redding, G. M. & Wallace, B. Calibration and alignment are separable: Evidence from prism adaptation. Journal of motor behavior 33, 401–412 (2001). URL https://doi.org/10.1080/00222890109601923.

91. OShea, J. et al. Kinematic markers dissociate error correction from sensorimotor realignment during prism adaptation. Neuropsychologia 55, 15–24 (2014). URL https://www.sciencedirect.com/science/article/pii/S0028393213003138.

92. Herwig, U., Satrapi, P. & Schönfeldt-Lecuona, C. Using the international 10-20 EEG system for positioning of transcranial magnetic stimulation. Brain Topography 16, 95–99 (2003). URL https://doi.org/10.1023/B:BRAT.0000006333.93597.9d.

93. Yousry, T. A. et al. Localization of the motor hand area to a knob on the precentral gyrus. A new landmark. Brain 120, 141–157 (1997). URL https://doi.org/10.1093/brain/120.1.141.

94. Engel, S. A., Glover, G. H. & Wandell, B. A. Retinotopic organization in human visual cortex and the spatial precision of functional MRI. Cerebral Cortex 7, 181–192 (1997). URL https://doi.org/10.1093/cercor/7.2.181.

95. Lunghi, C., Emir, U., Morrone, M. & Bridge, H. Short-term monocular deprivation alters gaba in the adult human visual cortex. Current Biology 25, 1496–1501 (2015). URL https://www.sciencedirect.com/science/article/pii/S0960982215004340.

96. Ip, I. B. et al. Combined fMRI-MRS acquires simultaneous glutamate and BOLD-fMRI signals in the human brain. NeuroImage 155, 113–119 (2017). URL https://www.sciencedirect.com/science/article/pii/S1053811917303233.

97. öz, G. & Tkač, I. Short-echo, single-shot, full-intensity proton magnetic resonance spectroscopy for neurochemical profiling at 4 T: Validation in the cerebellum and brainstem. Magnetic Resonance in Medicine 65, 901–910 (2011). URL https://onlinelibrary.wiley.com/doi/abs/10.1002/mrm.22708.

98. Deelchand, D. K. et al. Two-site reproducibility of cerebellar and brainstem neurochemical profiles with short-echo, single-voxel mrs at 3t. Magnetic Resonance in Medicine 73, 1718– 1725 (2015). URL https://onlinelibrary.wiley.com/doi/abs/10.1002/mrm.25295.

99. Natt, O., Bezkorovaynyy, V., Michaelis, T. & Frahm, J. Use of phased array coils for a determination of absolute metabolite concentrations. Magnetic Resonance in Medicine 53, 3–8 (2005). URL https://onlinelibrary.wiley.com/doi/abs/10.1002/mrm.20337.

100. Provencher, S. W. Estimation of metabolite concentrations from localized in vivo proton NMR spectra. Magnetic Resonance in Medicine 30, 672–679 (1993). URL https://onlinelibrary.wiley.com/doi/abs/10.1002/mrm.1910300604.

101. Provencher, S. W. Automatic quantitation of localized in vivo ^1^H spectra with lcmodel. NMR in Biomedicine 14, 260–264 (2001). URL https://onlinelibrary.wiley.com/doi/abs/10.1002/nbm.698.

102. Provencher, S. LCModel & LCMgui users manual. Stephen Provencher Inc (2012).

103. Porges, E. C., et al. Frontal gamma-aminobutyric acid concentrations are associated with cognitive performance in older adults. Biological Psychiatry: Cognitive Neuroscience and Neuroimaging 2, 38–44 (2017). URL https://www.sciencedirect.com/science/article/pii/S2451902216300696.

104. Good, C. D. et al. A voxel-based morphometric study of ageing in 465 normal adult human brains. NeuroImage 14, 21–36 (2001). URL https://www.sciencedirect.com/science/article/pii/S1053811901907864.

105. Harris, A. D., Puts, N. A. & Edden, R. A. Tissue correction for GABA-edited MRS: Considerations of voxel composition, tissue segmentation, and tissue relaxations. Journal of Magnetic Resonance Imaging 42, 1431–1440 (2015). URL https://onlinelibrary.wiley.com/doi/abs/10.1002/jmri.24903.

106. Porges, E. C. et al. Impact of tissue correction strategy on GABA-edited MRS findings. NeuroImage 162, 249–256 (2017). URL https://www.sciencedirect.com/science/article/pii/S1053811917307097.

107. Scholl, J. et al. Excitation and inhibition in anterior cingulate predict use of past experiences. eLife 6, e20365 (2017). URL https://doi.org/10.7554/eLife.20365.

108. Zhang, Y., Brady, M. & Smith, S. Segmentation of brain mr images through a hidden markov random field model and the expectation-maximization algorithm. IEEE Transa-tions on Medical Imaging 20, 45–57 (2001). URL https://ieeexplore.ieee.org/document/906424.

109. R Core Team. *R: A Language and Environment for Statistical Computing*. R Foundation for Statistical Computing, Vienna, Austria (2017). URL https://www.R-project.org/.

110. Hothorn, T., Bretz, F. & Westfall, P. Simultaneous inference in general parametric models. Biometrical Journal 50, 346–363 (2008).

111. Ben-Shachar, M. S., Makowski, D. & Ldecke, D. Compute and interpret indices of effect size. CRAN (2020). URL https://github.com/easystats/effectsize. R package.

112. Cohen, J. Statistical power analysis for the behavioral sciences (Academic press, 2013).

113. Stagg, C., Bachtiar, V. & Johansen-Berg, H. The role of GABA in human motor learning. Current Biology 21, 480–484 (2011). URL https://www.sciencedirect.com/science/article/pii/S0960982211001254.

114. Jocham, G., Hunt, L. T., Near, J. & Behrens, T. E. J. A mechanism for value-guided choice based on the excitation-inhibition balance in prefrontal cortex. Nature Neuroscience 15, 960–961 (2012). URL https://doi.org/10.1038/nn.3140.

115. Imai, K., Keele, L., Tingley, D. & Yamamoto, T. Causal mediation analysis using r. In Vinod, H. D. (ed.) Advances in Social Science Research Using R (Springer-Verlag, New York, 2010).

116. Cavassila, S., Deval, S., Huegen, C., Van Ormondt, D. & Graveron-Demilly, D. Cramr–rao bounds: an evaluation tool for quantitation. NMR in Biomedicine 14, 278–283 (2001).

117. Stone, J. M. et al. Ketamine effects on brain gaba and glutamate levels with 1h-mrs: relationship to ketamine-induced psychopathology. Molecular psychiatry 17 (2012).

118. Rowland, L. M. et al. In vivo measurements of glutamate, gaba, and naag in schizophrenia. Schizophrenia bulletin sbs092 (2012).

119. Tisell, A., Leinhard, O. D., Warntjes, J. B. M. & Lundberg, P. Procedure for quantitative 1h magnetic resonance spectroscopy and tissue characterization of human brain tissue based on the use of quantitative magnetic resonance imaging. Magnetic resonance in medicine 70, 905–915 (2013).

120. Kreis, R. The trouble with quality filtering based on relative cramér-rao lower bounds. Magnetic resonance in medicine 75, 15–18 (2016).

121. Minichiello, L. Trkb signalling pathways in LTP and learning. Nature Reviews Neuroscience 10, 850–860 (2009). URL https://doi.org/10.1038/nrn2738.

122. Kleim, J. A. et al. BDNF val66met polymorphism is associated with modified experience-dependent plasticity in human motor cortex. Nature Neuroscience 9, 735–737 (2006). URL https://doi.org/10.1038/nn1699.

123. McHughen, S. A., Pearson-Fuhrhop, K., Ngo, V. K. & Cramer, S. C. Intense training overcomes effects of the val66met bdnf polymorphism on short-term plasticity. Experimental Brain Research 213, 415 (2011). URL https://doi.org/10.1007/s00221-011-2791-z.

124. Joundi, R. A. et al. The effect of BDNF val66met polymorphism on visuomotor adaptation. Experimental Brain Research 223, 43–50 (2012). URL https://doi.org/10.1007/s00221-012-3239-9.

125. Chen, X. et al. A chemical-genetic approach to studying neurotrophin signaling. Neuron 46, 13–21 (2005). URL https://www.sciencedirect.com/science/article/pii/S0896627305002047.

126. Egan, M. F. et al. The BDNF val66met polymorphism affects activity-dependent secretion of bdnf and human memory and hippocampal function. Cell 112, 257–269 (2003). URL https://www.sciencedirect.com/science/article/pii/S0092867403000357.

127. Verhagen, M. et al. Meta-analysis of the BDNF val66met polymorphism in major depressive disorder: effects of gender and ethnicity. Molecular Psychiatry 15, 260–271 (2010). URL https://doi.org/10.1038/mp.2008.109.

